# Excised linear introns regulate growth in yeast

**DOI:** 10.1101/426049

**Authors:** Jeffrey T. Morgan, Gerald R. Fink, David P. Bartel

## Abstract

Spliceosomal introns are ubiquitous non-coding RNAs typically destined for rapid debranching and degradation. Here, we describe 34 excised *Saccharomyces cerevisiae* introns that, although rapidly degraded in log-phase growth, accumulate as linear RNAs under either saturated-growth conditions or other stresses that cause prolonged inhibition of TORC1, a key integrator of growth signaling. Introns that become stabilized remain associated with components of the spliceosome and differ from the other spliceosomal introns in having a short distance between their lariat branch point and 3′ splice site, which is necessary and sufficient for their stabilization. Deletion of these unusual introns is disadvantageous in saturated conditions and causes aberrantly high growth rates of yeast chronically challenged with the TORC1 inhibitor rapamycin. Reintroduction of native or engineered stable introns suppresses this aberrant rapamycin response. Thus, excised introns function within the TOR growth-signaling network of *S. cerevisiae*, and more generally, excised spliceosomal introns can have biological functions.

---

Spliceosomal introns are a defining feature of eukaryotic life; they are present in all known eukaryotic genomes and absent from all known non-eukaryotic genomes^1,2^. Every splicing event produces two products: ligated exons and an excised lariat intron^3-7^. Because their production is an obligate result of gene expression and mRNA maturation, introns could be a fertile source of functional ncRNAs across eukaryota. However, although produced abundantly, excised lariat introns are debranched and degraded within seconds^8-11^. So, although introns have broad roles in essential alternative splicing events during pre-mRNA processing^12^, excised introns are generally viewed not as products of splicing but instead as inactive byproducts of exon ligation^13^.

Although not reported to accumulate post-splicing, introns can play important roles either before or after splicing. Function for individual introns has been examined most thoroughly in the intron-poor budding yeast *S. cerevisae*, which contains approximately 300 spliceosomal introns, with only 14 multi-intronic genes and only a few annotated cases of alternative-splicing events^14-17^. Thus, in yeast, potential functions of introns after splicing can be more readily separated from their functions during pre-mRNA processing. For most introns tested, no growth phenotypes are detected upon intron removal^18-20^. In a few cases, however, functions have been observed. These functions include regulating expression of duplicated ribosomal protein genes^21^ and counteracting R-loop formation during transcription^22^. In these cases, the function manifests entirely during pre-mRNA production and processing and thus before the intron exists as a separate RNA molecule. With respect to functions post-splicing, some introns are processed to produce noncoding RNAs, such as small nucleolar RNAs (snoRNAs), although in these cases the flanking portions of the intron are still rapidly catabolized^23,24^. Thus, functional analyses in *S. cerevisiae* support the prevailing view that the collective fate of introns post-splicing is solely to be debranched and at least partially degraded.

Although hundreds of individual yeast introns have been assayed for function, relatively few experimental conditions have been explored. Most experiments assay cells in the exponential growth phase, which provides consistent measurements in a standardized system sensitized to detect differences in growth and metabolism. However, outside of the laboratory setting, yeast cells are unlikely to spend many consecutive generations rapidly dividing and more often face limiting nutrients or other stresses^25^. Because the ability of cells to appropriately respond to these suboptimal conditions would have been important for survival, the findings from more optimal environments might not reflect biological phenomena present during non-exponential phases of growth that more adequately reflect growth in natural habitats. Accordingly, we set out to examine gene regulation of *S. cerevisiae* outside of the context of log-phase growth.

## Accumulation of excised, linear introns

We performed RNA sequencing (RNA-seq) on two *S. cerevisiae* samples: one taken from a culture in log-phase growth and the other from a saturated culture in which cell density was minimally increasing. For most intron-containing genes, such as that of actin *(ACT1*), very few intron-mapping RNA-seq reads were observed from either culture condition (Fig. 1a), as expected if introns were rapidly degraded post-splicing^9^. However, for a subset of genes, exemplified by *ECM33*, many intron-mapping reads were observed specifically from the saturated culture (Fig. 1b). Indeed, the much higher density of reads for the *ECM33* intron compared to that for its exons suggested that in this growth condition, the intron accumulate *ECM33* intron, coverage dropped abruptly at the 5′ and 3′ boundaries of the intron (Fig. 1b), indicating the intron was not being retained in the mature mRNA.

**Figure 1.**
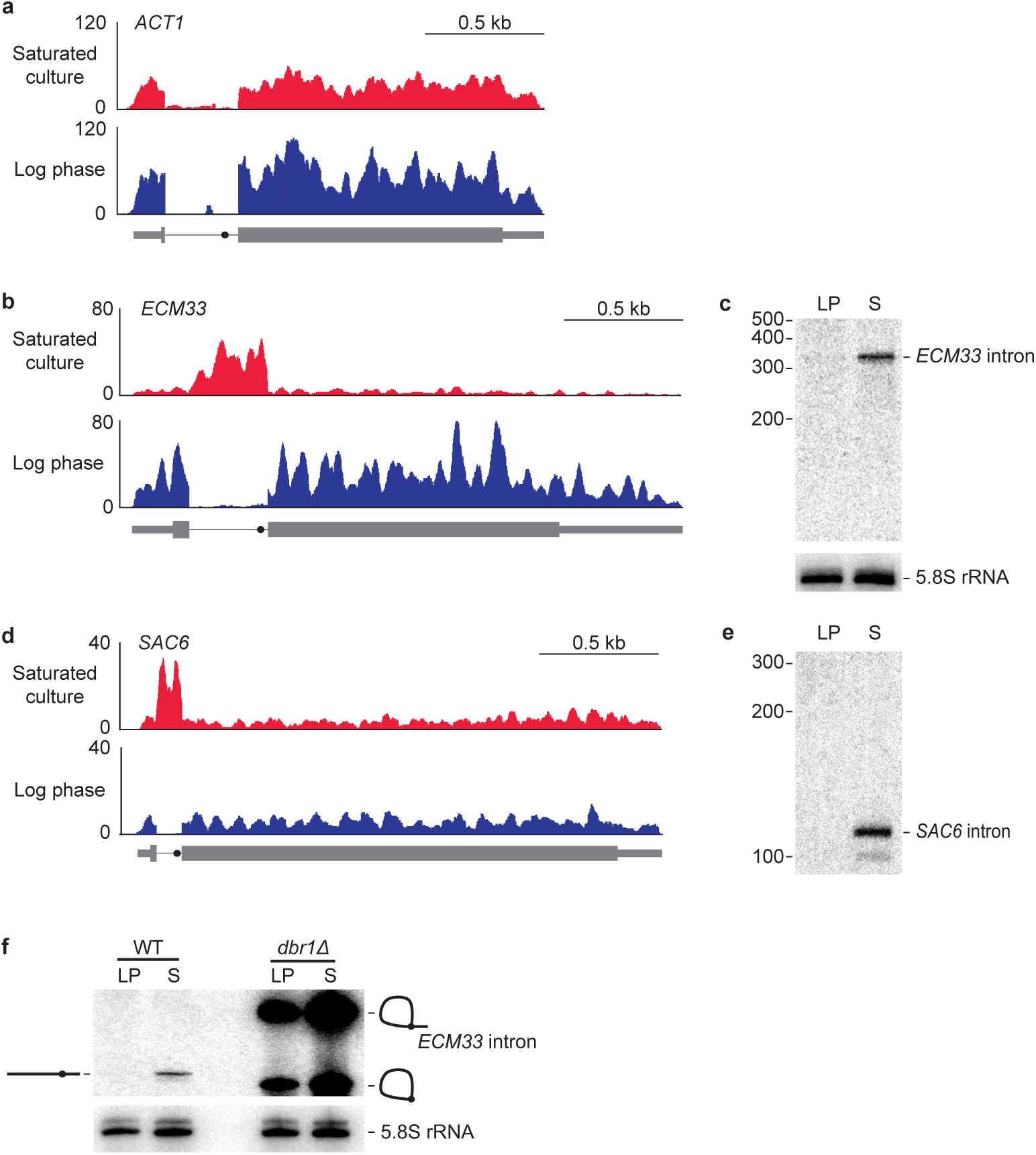
Some excised linear introns accumulate in yeast. **a**, Undetectable accumulation of the intron from the *ACT1* gene. Shown are RNA-seq profiles for *ACT1* (thick box, CDS; thin box, UTR; line, intron; closed circle on intron, BP) in both log-phase and saturated yeast cultures. Plotted for each nucleotide of the locus is the number of reads per million combined exon- and intron-mapping reads of the library. **b**, Accumulation of the *ECM33* intron in saturated but not log-phase culture, otherwise as in **a**. **c**, Accumulation of intact *ECM33* intron in saturated but not log-phase culture. Shown is an RNA blot that resolved total RNA from both log-phase (LP) and saturated (S) cultures and was probed for the *ECM33* intron. Migration of markers with lengths indicated (nucleotides) is at the left. For a loading control, the blot was reprobed for 5.8S rRNA. **d**, Accumulation of the *SAC6* intron in saturated but not log-phase culture, otherwise as in **a**. **e**, Reprobing of RNA blot in **c** for the *SAC6* intron; otherwise, as in **c**. **f**, Accumulation of the debranched form of the *ECM33* intron in a saturated culture. Analysis is as in **c**, except the RNA blot included samples from both WT and *dbr1Δ* cultures. Migration of the linear isoform is indicated (diagram on the left) as is migration of both the lariat and the circle isoforms (diagrams on the right).

To examine whether our RNA-seq analysis was indeed detecting accumulation of excised introns of a defined size, we probed RNA blots for the inferred RNA species. Probes to the *ECM33* intron detected a single major species running at the position expected for the full-length, 330-nt intron (Fig. 1c). Likewise, a probe for *SAC6*, another intron inferred by RNA-seq to accumulate in the saturated culture (Fig. 1d), detected a defined RNA of the size expected for the full-length excised intron (Fig. 1e).

A known mechanism by which excised introns can be protected from degradation is if they evade debranching and persist as either lariat RNAs or circular lariat derivatives in which the lariat tail is missing^10,26,27^. However, these nonlinear species have either branched RNA or a 2’–5’ phosphodiester linkage, which would impede reverse transcriptase^5,28^ and thereby cause RNA-seq reads to be depleted in the region of the branch point—a pattern that we did not observe in the RNA-seq profiles (Fig. 1b, Extended Data Fig. 1). To test further whether *ECM33* intronic RNA accumulated as either a lariat RNA or its circular derivative, we harvested RNA from yeast lacking Dbr1, the enzyme required to debranch intron lariats^10^, and compared the *ECM33* intronic RNA that accumulated in the *dbr1Δ* strain with RNA that accumulated in wild-type saturated cultures. As expected, in *dbr1Δ* log-phase culture, *ECM33* intron was detected as two abundant species, which corresponded to the branched lariat and circle, and the same species accumulated in *dbr1Δ* saturated culture (Fig. 1f). Importantly, neither of these nonlinear species co-migrated with the linear intron identified in wild-type saturated culture. These results confirmed that Dbr1 is necessary to form linear introns outside of log phase and showed that the previously known mode of decreased intron turnover cannot explain the observed accumulation of excised introns in a saturated *S. cerevisiae* culture.

Another mechanism that might protect these introns from degradation in saturated cultures is incorporation into a ribonucleoprotein complex. Indeed, the *ECM33* intron predominantly co-sedimented with complexes about the size of ribosomal subunits (35–50 Svedberg units, Extended Data Fig. 2a). To identify proteins associating with the intron, we performed pull-down experiments from gradient fractions containing the intron and used quantitative mass spectrometry to identify the co-purifying proteins. For these experiments, we took advantage of the observation that tagged versions of the *ECM33* intron excised from expression constructs retained the behavior of the endogenous *ECM33* intron, i.e., rapid degradation in log-phase culture and accumulation as excised linear RNA in saturated culture (Extended Data Fig. 2b, c). The top 10 proteins consistently copurifying with MS2-tagged versions of the *ECM33* intron (average enrichment = 4.9 fold) were each spliceosomal proteins, the identities of which indicated that the excised and debranched *ECM33* intron resided in a specific complex, which when compared to known spliceosome complexes most closely resembled the intron-lariat spliceosome (ILS) complex (Extended Data Table 1)^29^.

Studies in log-phase extracts indicate that introns are debranched after spliceosome disassembly^30^. If stable introns were also debranched outside the spliceosome, they would then need to re-associate with spliceosome components as linear RNAs. A simpler model is of continued association of the linear RNAs with ILS components post-splicing, in which case the relationship between ILS disassembly and debranching presumably varies, depending on growth condition and/or intron identity. In either scenario, our findings indicate that in saturated cultures accumulating introns are bound to and presumably protected by a complex resembling the ILS.

## Defining features of stable introns

We performed a systematic search for all introns that undergo a switch in stability and accumulate as linear RNAs in saturated cultures, hereafter referred to as “stable introns.” In this search, RNA-seq reads of each intron-containing gene were analyzed for a preponderance of reads mapping to introns, particularly those mapping precisely the edges of the excised introns (consistent with a post-splicing intron) relative to those mapping across splice sites and splice junctions (signatures of intron retention and mature mRNA expression, respectively)^31^ (Fig. 2a). Inspection of RNA-seq reads that mapped to the 3′ edges of stable introns identified many that were extended by one or more untemplated adenosine residues (Fig. 2b). This frequent addition of untemplated adenosines was not observed on either reads with 3′ ends mapping to the interior of stable introns (Fig. 2b) or reads mapping to introns in log phase—although very few reads were obtained for this latter case. Short 3′-terminal oligo(A) tails are added by TRAMP complex to mark nuclear RNAs for degradation^32,33^ and have been observed on lariat introns isolated from *dbr1Δ* yeast^34^. Our finding that many stable-intron molecules had these tails suggests that these molecules might have been targeted for exosomal decay yet were somehow protected from this decay. Regardless of their function, these untemplated adenosine residues provided an additional criterion for the annotation of stable introns, which helped us confidently identify another 28 introns whose stable form accumulated in yeast grown in saturated cultures (Extended Data Fig. 3, Extended Data Table 2).

**Figure 2.**
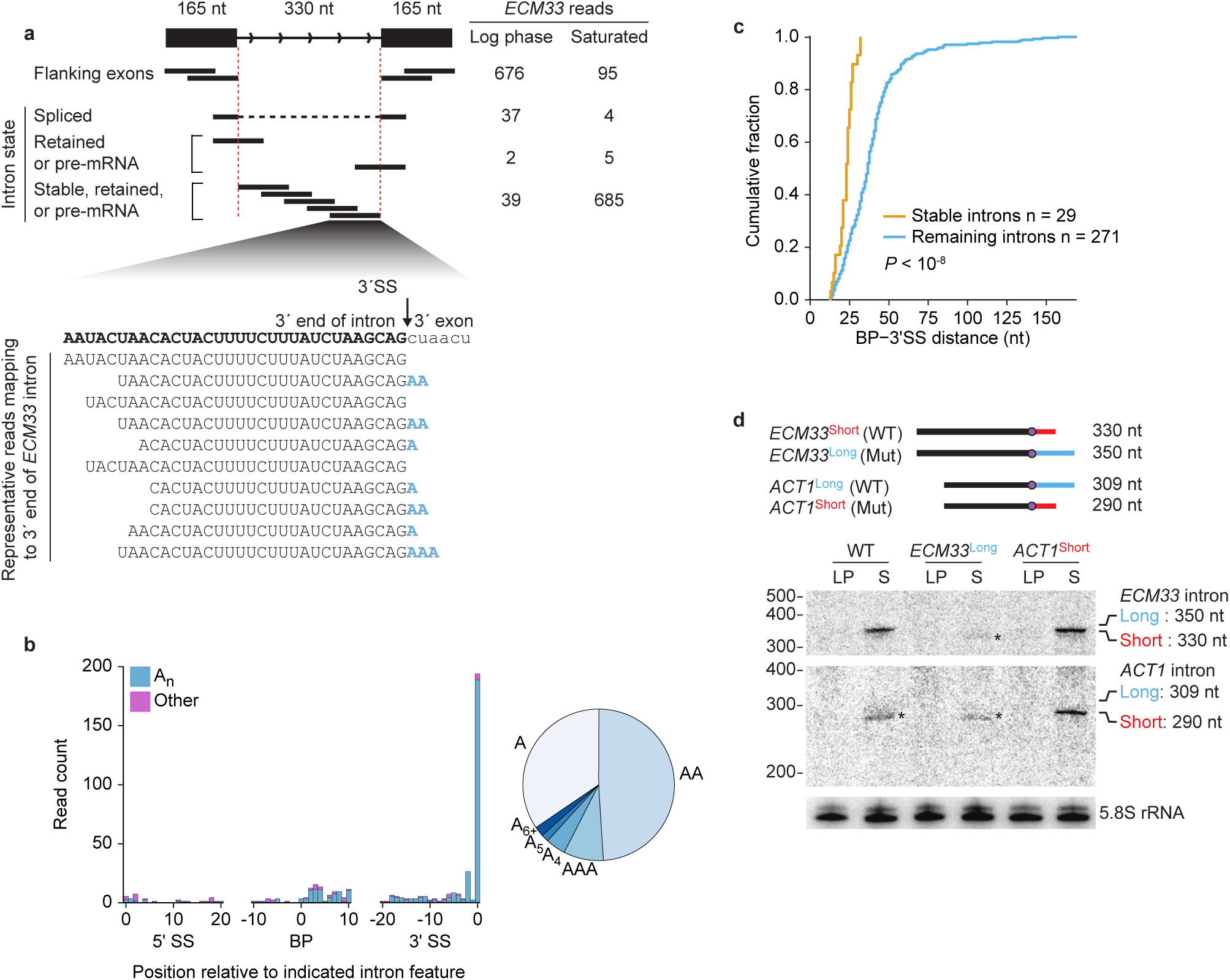
Stable introns have oligo(A) tails and short BP–3′SS distances. **a**, Example of RNA-seq support for intron accumulation and terminal adenylylation. The diagram (top left) classifies the RNA-seq reads deriving from the possible intron states of *ECM33* transcripts (black lines, reads; red dashed lines, intron boundaries). Read counts from logphase and saturated cultures (normalized for library depth) are listed for each class of reads (top right). For convenient comparison to intron accumulation, exon reads are only counted if they map within 165 nucleotides of either splice site (thereby encompassing 330 nucleotides, the length of the intron). The alignment (bottom) shows representative reads mapping to the intron 3′ terminus, aligned below the sequence of the 3′ intron-exon junction. Many of these reads had untemplated terminal adenosine residues (blue). **b**, Composition of untemplated tailing nucleotides observed in a saturated culture. Reads that still had at least one terminal untemplated nucleotide after trimming the 3′-adapter sequence were collected, and the position of this tail was annotated as that of the last templated nucleotide. Counts of reads with tails added at positions 0 to +20 relative to the 5′SS, –10 to +10 relative to the BP, and –20 to 0 relative to the 3′SS are plotted, binning counts for tails of only adenosines (An, teal) separately from those of all other tails (other, purple). For An tails mapping to the 3′-terminal nucleotide of introns, the fraction with each indicated length is plotted (right). The relative abundance of A_n_ at positions –2 and –1 relative to the 3′SS was ambiguous because most introns have an A at position –1 (3′SS consensus sequence, YAG), which causes tails of length N at position –1 to be indistinguishable from tails of length N+1 at position –2. **c**, The shorter BP–3′SS distance of stable introns compared to that of most other introns. Plotted are cumulative distributions of BP–3′SS distances (*P* < 10^-8^, one–tailed Kolmogorov-Smirnov test). **d**, A causal relationship between BP–3′SS distance and stable-intron formation. The diagram at the top shows WT and mutant (Mut) introns, in which the WT *ECM33* BP–3′SS distance was extended from 25 nt (short, red) to 45 nt (long, blue), and the WT *ACT1* BP–3′SS distance was shortened from 44 nt (long, blue) to 25 nt (short, red). Below the diagram are results of an RNA blot that resolved samples from the indicated strains and was probed sequentially for the *ECM33* intron (top), the *ACT1* intron (middle), and 5.8S rRNA. Probes were complementary to portions of introns common to both short and long isoforms. Migration of markers with lengths indicated (nucleotides) is at the left. Expected migration of long and short linear isoforms of each intron is at the right. The asterisks (∗) mark the detection of long-isoform degradation products, which each migrated even faster than did the short isoform.

**Figure 3.**
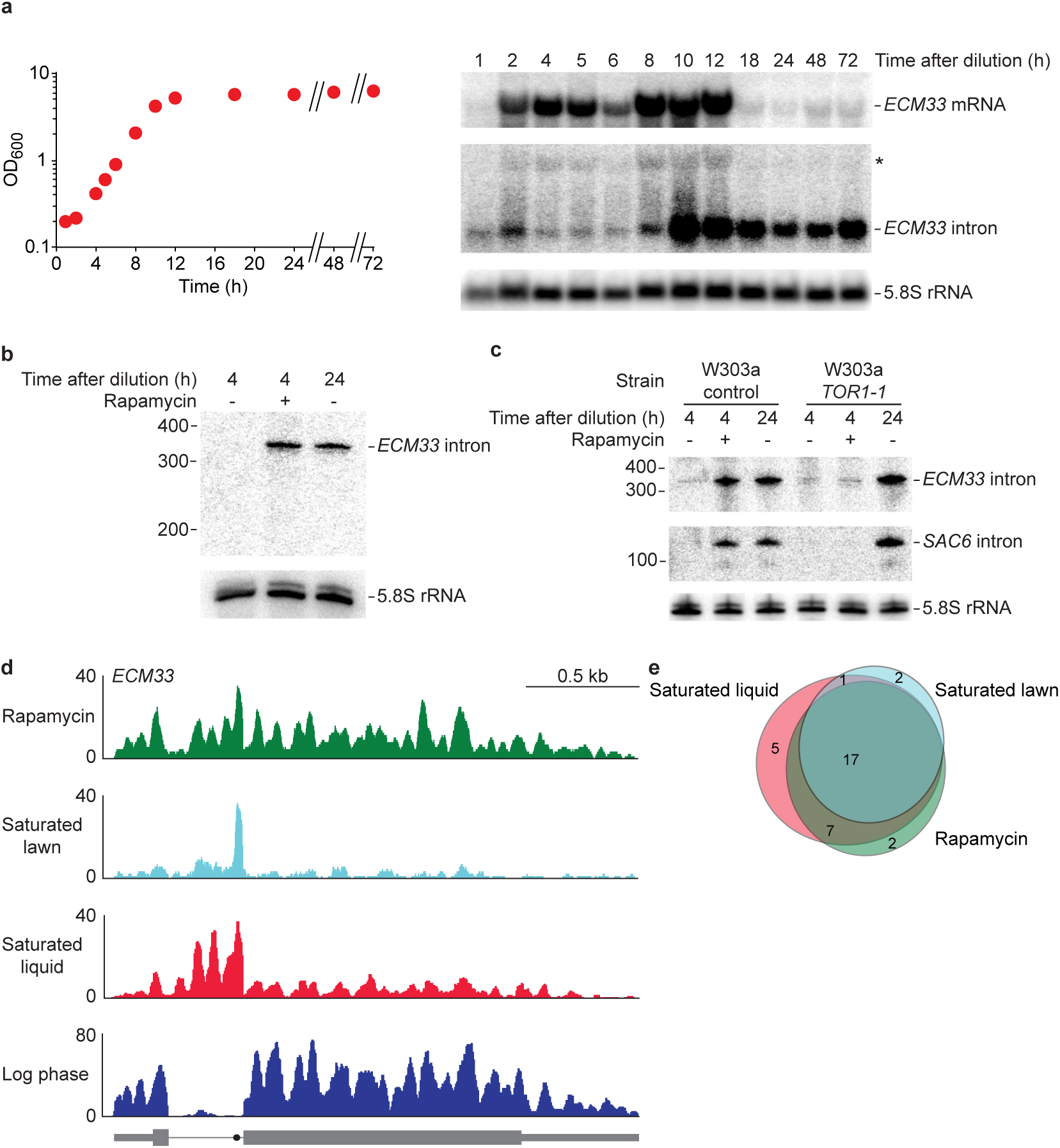
TORC1 regulates stable-intron formation. **a**, The distinct accumulation pattern of the *ECM33* stable intron and its host mRNA. Culture was seeded at OD_600_ (optical density at 600 nm wavelength) = 0.2 from an overnight culture and at the indicated time points growth was monitored using OD_600_ measurements (left), and aliquots were harvested for RNA-blot analysis (right). The RNA blot was as in Fig 1c, except it utilized a gel designed to resolve longer RNAs. The asterisk (∗) shows the migration of the *ECM33* pre-mRNA, which is also detected by the probe for the *ECM33* intron. **b**, Induced accumulation of the *ECM33* stable intron in cells cultured in the presence of rapamycin. To prevent contamination of starting cultures with stable introns contributed by inoculum, cultures were seeded at OD_600_ = 0.2 from an overnight culture, allowed to grow to early log phase, collected by vacuum filtration, and resuspended in fresh media without (lanes 1 and 3) or with (lane 2) 100 nM rapamycin. Cultures were harvested after indicated times and analyzed as in Fig. 1c. **c**, Requirement of rapamycin-sensitive *TORC1* for induction of stable introns by rapamycin. Compared are the results of a W303a control strain and a W303a strain containing a rapamycin-insensitive allele of *TOR1* (*TOR1-1* [S1972I])^56^. Otherwise, this panel is as in **b**, except RNA blot was additionally reprobed for the *SAC6* intron. **d**, Accumulation of the *ECM33* intron in rapamycin-treated, saturated-lawn, and saturated-liquid cultures, detected using RNA-seq. Results showing accumulation of this intron in saturated-liquid but not log-phase liquid culture are from a different biological replicate (samples from **b**, lanes 1 and 3) than those shown in Fig. 1a and b; otherwise, this panel is as in Fig. 1a. **e**, Overlap between stable introns identified in saturated-liquid, saturated-lawn, and rapamycin-treated cultures.

In several cases, one of these stable introns derived from one of the few yeast genes with more than one intron (e.g., *EFM5*, Extended Data Fig. 3). The differential stability of one intron but not of the other intron from the same gene suggested that intron-intrinsic characteristics drive stability and accumulation. Consequently, we searched for common features among the 30 stable introns, which the cellular machinery might use to differentiate stable introns from the majority of introns that are still rapidly degraded in saturated cultures. This search found that stable introns were indistinguishable from other introns in nearly every respect. Compared to other introns, they had similar strengths of canonical splicing motifs (Extended Data Fig. 4a), similar length distributions (Extended Data Fig. 4b, *P* > 0.05), no common predicted structures or enriched sequence motifs (Extended Data Fig. 4c and d), and no enriched functional ontologies of their host genes. Of the features examined, the only one that differed was the distance between the lariat branch point (BP) and 3′ splice site (3′SS), which tended to be shorter for stable introns (Fig. 2c, *P* < 10 ^-8^).

To investigate a potential role of BP position in influencing intron stability, we made mutations that changed endogenous BP-3′SS distances and examined the effects of these mutations on intron accumulation. Lengthening the short BP-3′SS distance of the normally stable *ECM33* intron from 25 to 45 nt abrogated accumulation of the full-length excised intron, indicating that a short BP–3′SS distance is required for stability of this intron (Fig. 2d, compare *ECM33*^short^ and *ECM33*^long^). Moreover, shortening the BP–3′SS distance of the normally unstable *ACT1* intron from 44 to 25 nt conferred stability to this intron in a saturated culture, which suggested that a short BP–3′SS distance is not only necessary but also sufficient for the stability of introns in saturated cultures (Fig. 2d, compare *ACT1*^short^ and *ACT1*^long^).

The notion that a short BP–3′SS distance is sufficient for stability seemed at odds with the observation that some introns with BP–3′SS distances of 20–25 nt were not annotated as stable introns (Fig. 2c). One possibility was that some introns with short BP–3′SS distances were not annotated as stable introns simply because their genes were not actively transcribed during the period at which stable introns were protected from degradation. To investigate this possibility, we placed introns that had not been identified as stable introns into the previously used expression construct, choosing two introns with a short (20- and 25-nucleotide) and two with a long (37- and 44-nucleotide) BP–3′SS distance. The two test introns with short BP–3′SS distances accumulated specifically in saturated cultures, whereas the two test introns with long BP–3′SS distances were unstable in both conditions (Extended Data Fig. 4e).

We conclude that the two defining features of stable introns are 1) a short BP–3′SS distance and 2) expression within a cellular context in which introns are stabilized. After being modified to satisfy these criteria, all tested non-stable introns became stable introns (Fig. 2d, Extended Data Fig. 4e), which suggested that no other sequence or structural features of either the intron or the host gene are required to achieve stability.

**Figure 4.**
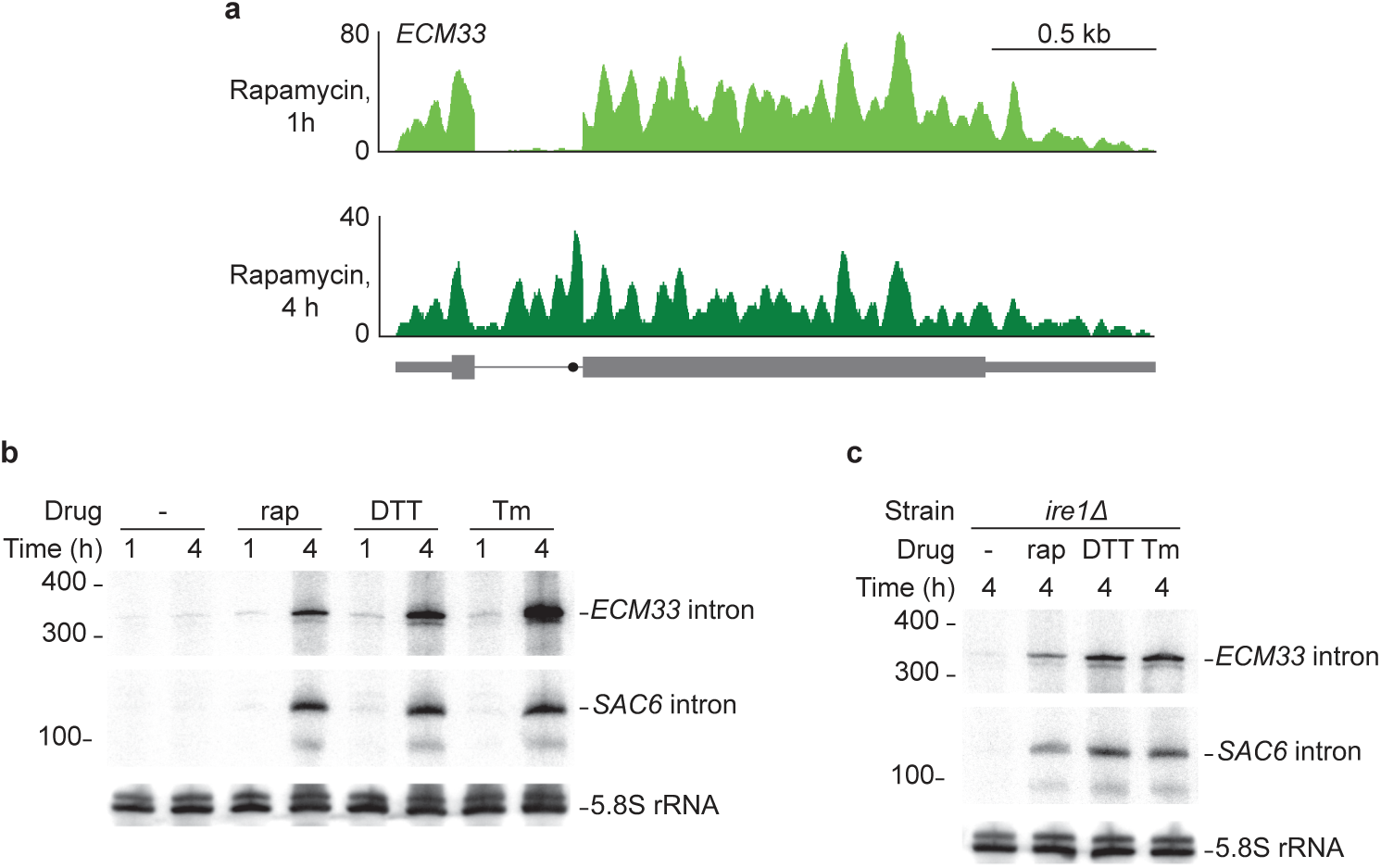
Stable-intron accumulation occurs after prolonged inhibition of TORC1. **a**, Undetectable accumulation of the *ECM33* intron after 1 h of rapamycin treatment. For comparison, results showing accumulation of this intron after 4 h of rapamycin treatment are re-plotted from Fig. 3c. Otherwise, this panel is as in Fig. 3c. **b**, Induced accumulation of stable introns in cells cultured with 5 mM DTT or 1 μg mL^-1^ tunicamycin. Otherwise, this panel is as in Fig. 3b. **c**, Accumulation of stable introns in cells cultured with DTT or tunicamycin despite deletion of *IRE1*. Otherwise, this panel is as in **b**.

## Regulation of stable introns

We next examined when, during the interval between log-phase and saturated conditions, the switch in intron stability occurs. This was accomplished by harvesting samples through 72 h of culture and monitored the levels of the *ECM33* stable intron and its host mRNA (Fig. 3a). The *ECM33* mRNA increased 2 h after culture seeding, remained steady through 12 h, and then decreased to low levels through the remainder of the time course (Fig. 3a). The Ecm33 protein is a cell-wall–related protein^35^, which explained its high expression during phases of rapid cell division. The *ECM33* intron had a very different pattern of accumulation. Intron levels were low through 8 h of growth, and then, as cells exited the rapid-growth phase, intron levels dramatically increased (Fig. 3a). Intron abundance remained high for at least the next 62 h, even as the levels of mature mRNA decreased 23 fold. Although the contrasting dynamics of the *ECM33* intron and mRNA illustrated the different behaviors sometimes observed for stable introns and their host mRNAs, the large decrease in the levels of the mature mRNA in a saturated culture was not observed for all mRNAs that hosted stable introns. Indeed, as a class, these mRNAs had no significant trend in expression between log-phase and saturated cultures (Extended Data Fig. 4f).

The timing of stable-intron accumulation suggested that their formation might be linked to exit from rapid growth. With this in mind, we inhibited the target of rapamycin complex 1 (TORC1), a broadly conserved master integrator of nutritional and other environmental signals^36,37^, using the small molecule rapamycin. In yeast, rapamycin-mediated TORC1 inhibition leads to repression of anabolic processes, stimulation of catabolic processes, and ultimately a greatly reduced growth rate^38^. When a log-phase culture was resuspended in fresh media containing rapamycin, *ECM33* intron accumulated after 4 h (Fig. 3b). A repeat of this experiment using a strain with a rapamycin-resistant allele of *TOR1* (*TOR1-1*) yielded no accelerated intron accumulation upon treatment with rapamycin (Fig. 3c). *TOR1-1* cultures nonetheless accumulated stable introns upon reaching saturation (Fig. 3c), as expected in this strain known to be sensitive to endogenous TORC1 inhibition^39^. Thus, rapamycin-induced intron accumulation was specifically due to inhibition of TORC1. Inhibition of protein synthesis in a TORC1-independent way did not cause stable-intron formation (Extended Data Fig. 5a). Moreover, stable introns accumulated to lower levels in TORC1^∗^, a strain harboring six hyperactive alleles of the regulatory network^40^ (Extended Data Fig. 6), a result that further validated the involvement of TORC1 signaling.

To extend this investigation transcriptome-wide, we prepared RNA-seq libraries from yeast treated for 4 h with rapamycin. We also investigated stable-intron formation in a metabolically different^41^—but experimentally common—saturated-growth scenario: a lawn of yeast grown over 3 days in an aerobic environment. When compared to the saturated liquid culture, both rapamycin-treated cells and cells from the lawn showed similar accumulation not only of the *ECM33* intron (Fig. 3d) but also of other stable introns (Fig. 3e). In total, 34 introns (11% of *S. cerevisiae* introns) were classified as stable introns in at least one of the three conditions (Extended Data Fig. 3, Extended Data Table 2). A somewhat greater yield was obtained from the saturated liquid culture (Fig. 3e), presumably because deeper sequencing of this sample enabled confident identification of more lowly expressed stable introns, implying that with even deeper sequencing, additional stable introns would be confidently identified.

We tested several genetic and environmental perturbations for their effects on stable-intron accumulation. Gain-of-function mutations in *TAP42* or *SCH9* (the two major effector branches of TORC1) were individually insufficient to override TORC1 repression and attenuate stable-intron formation (Extended Data Fig. 5b,c), and Sch9 activity was not required for stable-intron regulation (Extended Data Fig. 5d), which implicated involvement of another effector branch of TORC1. Although depletion of carbon, nitrogen, and amino acids rapidly inhibit facets of TORC1 signaling^36^, these conditions did not induce pre-mature stable-intron formation (Extended Data Fig. 5e). Likewise, direct perturbations of the general amino acid control [GAAC] pathway did not disrupt stable-intron regulation (Extended Data Fig. 5f,g). Thus, the role of TORC1 in stable-intron regulation appeared separable from its role in the rapid response to nutrient deprivation.

As illustrated for the *ECM33* intron (Fig. 4a), stable introns were not detected after treating with rapamycin for 1 h, a time period sufficient to phenocopy the TORC1 response to nutrient deprivation. This result agreed with our observation that some conditions known to rapidly inhibit aspects of TORC1 signaling did not induce stable introns and led us to explore the time sensitivity of response to rapamycin. Few studies have explored prolonged TORC1 inhibition in yeast, and those that report on ≥2 h rapamycin treatments observe strikingly different TORC1-mediated responses compared to short-term rapamycin treatment or starvation^42-45^. Interestingly, some of these studies^43-45^ show that prolonged TORC1 inhibition (≥2 h treatment with rapamycin or tunicamycin) phenocopies secretory stress more closely than nutrient stress. Indeed, we found that the secretory stressors tunicamycin and DTT induced stable-intron formation, that this induction required similar, extended treatment durations as found with rapamycin, and that induction of stable introns by these canonical unfolded-protein response (UPR) activators did not require the essential UPR sensor *IRE1* (Fig. 4b,c). These results indicated that TORC1-mediated stable-intron accumulation depends on connections between TORC1 and membrane-trafficking/secretory stress^46,47^, and not its more oft-considered connections to nutrient stress.

## Biological function of stable introns

The discovery of stable introns and their link to the TOR growth-signaling pathway brought to the fore the question of their function. To test for loss-of-function phenotypes, we used a CRISPR-Cas9 system adapted for *S. cerevisiae*^48,49^ to precisely remove introns without affecting exonic sequences. Reasoning that stable introns might contribute collectively to function, and thus a phenotype might be detected only after multiple introns were deleted from the same strain, we generated a quintuple mutant lacking stable introns of five genes (*ecm33, ubc4, hnt1, sac6*, and *rfa2*). Because intron from these genes normally accumulated to high levels, this strain, called EUHSR, had less than half the stable intron molecules as the wild-type (WT) strain (Fig. 5a, Extended data Fig. 7). To detect the consequences of reducing stable-intron abundance, we co-cultured the EUHSR and wild-type strains in a competitive-growth assay. The co-culture was grown for three days to confluence, diluted, grown for three days to confluence again, and so on for a total of six cycles of growth. Time points taken just before dilution and after a day of regrowth revealed a saw-tooth pattern in which the WT strain had a small but significant advantage during the period of sustained saturated culture, whereas the EUHSR had an even greater advantage during the period that included re-entry to growth and exit from growth (Fig. 5b). These results indicated that stable-intron expression is beneficial in some physiological contexts and detrimental in others.

**Figure 5.**
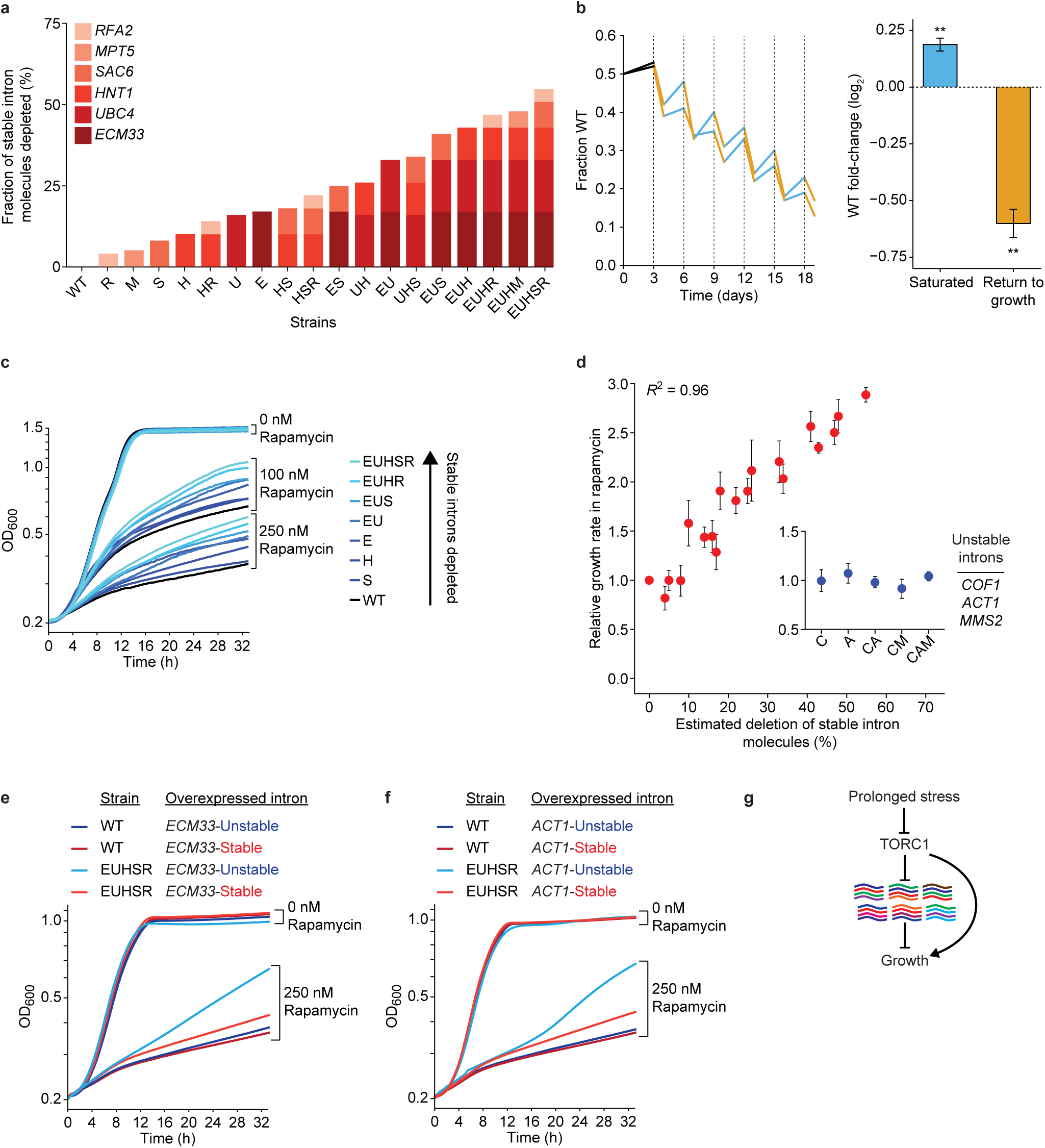
Stable introns influence yeast fitness and growth. **a**, Eighteen strains with genomic deletions that deplete stable-intron molecules. In each strain, introns from 1–5 of the genes listed in the key were deleted. Deletion strains are named using single-letter abbreviations of the genes with deleted introns. The bar for each strain plots its estimated depletion of stable-intron molecules, indicating the contribution of each deleted intron (color key) based on its contribution to the total number of stable-intron molecules in WT saturated culture. **b**, Altered fitness of the EUHSR strain, which had fewer stable-intron molecules. Left, results of competitive growth, in which the WT and EUHSR were mixed and then co-cultured, diluting the co-culture every three days (dashed lines). Plotted is the fraction of WT in the culture, measured twice for each cycle of growth, once just before dilution and again after a day regrowth. Lines for each of the two biological replicates are segmented to indicate periods of saturated growth (blue), periods of returning to growth (orange), and the initial post-seeding segment (black, which was not considered for calculations on the right). Right, mean fold-change of WT in either the periods of saturated growth (blue) or periods of return to growth (orange) (error bars, s.e.m.). Both mean fold-changes were statistically significant (∗∗, P <10^-4^, unpaired two-tailed t test). **c**, Attenuated rapamycin response of strains with fewer stable-intron molecules. Shown are growth curves of the indicated deletion strains (color key) when cultured in the presence of either 0 nM, 100 nM, or 250 nM rapamycin. Each curve shows the average for technical triplicates from a representative 96-well plate assaying eight strains. **d**, Relationship between the depletion of stable-intron molecules and increased growth in 250 nM rapamycin. Estimated depletion of stable-intron molecules is as in **b**. Relative growth rates are averages of biological replicates (n = 12 for WT, n = 3 for all other strains; error bars, s.e.m.). Also shown are results for control strains in which 1–3 unstable introns were deleted in the indicated genes (inset). **e**, Rescue of the rapamycin-response defect by ectopically expressing the *ECM33* stable intron. Shown are representative growth curves of WT and EUHSR strains overexpressing either the native stable *ECM33* intron (red) or an unstable *ECM33* intron (blue) cultured in the presence of either 0 nM or 250 nM rapamycin. The unstable version of the intron (*EMC33*-Unstable) is described in Fig. 3d (*EMC33*^Long^). Each curve plots the averages of technical triplicates. **f**, Rescue by an engineered *ACT1* stable intron. As in **e**, but ectopically expressing either the native unstable *ACT1* intron (blue) or an engineered stable *ACT1* intron (red). The stable version of the intron (*ACT1*-Stable) is described in Fig. 2d (*ACT1*^Short^). **g.** Placement of stable introns in the TORC1-mediated environmental-response pathway. TORC1 represses accumulation of diverse stable introns (colored lines), which in turn repress cell growth. Prolonged stress from diverse sources (rapamycin, tunicamycin, and DTT) inhibits the ability of TORC1 to repress stable-intron accumulation. This repressor of a repressor leads to growth stimulation, as do previously characterized branches of the pathway, which do not involve stable introns (arrow bypassing stable introns).

Because of the connection between TORC1 inhibition and stable-intron formation (Fig. 3b–e), we tested the effect of rapamycin on growth of mutant strains from an allelic series of single, double, triple, quadruple, and quintuple intron deletions that produced decreasing amounts of stable introns, culminating with the EUHSR quintuple mutant (Fig. 5a). In the absence of rapamycin, the strains all grew at equivalent rates (Fig. 5c). In the presence of rapamycin, growth was inhibited for all strains, but this growth inhibition was attenuated for strains lacking one or more stable intron, with mutant strains lacking more stable introns growing at faster rates (Fig. 5c). Indeed, across all the strains assayed, we observed a striking correlation between the estimated fraction of stable-intron molecules depleted from the transcriptome and attenuation of the rapamycin response (Fig. 5d, Pearson *R*^2^ = 0.96). In contrast, the rapamycin response of control strains that lacked nonstable introns of highly expressed genes was indistinguishable from that of wild type (Fig. 5d, inset), which showed that the observed phenotype was specific to depletion of stable introns and not a generic consequence of decreased splicing load.

The attenuated response to TORC1 inhibition seemed to depend solely on the number of stable-intron molecules that were removed from the cell and not on the identities of either the removed introns or their mutated host genes (Fig. 5d). This result provided evidence against the idea that the phenotype we observed might have been the result of subtle changes in host-gene expression or other secondary effects of intron removal or strain construction (such as off-target effects of the gene editing).

To further test the conclusion that the attenuated response to TORC1 inhibition depended solely on the aggregate number of stable-intron molecules produced and not on other factors, we examined the ability of ectopically expressed introns to rescue the phenotype. We first overexpressed the *ECM33* stable intron or, as a control, its unstable derivative that had a lengthened BP–3′SS distance (*ECM33*-stable and *ECM33*-unstable, respectively). The stability of this overexpressed intron had no detectable effect on either WT or EUHSR growth in the absence of rapamycin, and it had little effect on growth of the WT strain in rapamycin (Fig. 5e). Importantly, however, the stability of this ectopically expressed intron dramatically influenced the growth of the quintuple mutant in the presence of rapamycin—nearly completely rescuing its defective response to rapamycin (Fig. 5e). Analogous results were observed when ectopically expressing the stabilized and normal version of the *ACT1* intron (*ACT1*-stable and *ACT1*-unstable, respectively) (Fig. 5f), The ability of a stable version of the *ACT1* intron, which does not exist naturally in *S. cerevisiae*, to rescue the quintuple-mutant phenotype confirmed that this phenotype was a primary consequence of reduced stable-intron abundance.

Taken together, our results show that stable introns function within TORC1-mediated stress response in *S. cerevisiae* (Fig. 5g). TORC1 activity prevents stable-intron formation, as shown by the accumulation of these introns when cells undergo prolonged TORC1 inhibition (Fig. 3b–e, Fig. 4), and the stable introns inhibit growth of cells with decreased TORC1 activity, as shown by the increased growth observed in TORC1-inhibited strains with fewer stable introns (Fig. 5). Thus, this double-negative regulation, in which TORC1 inhibits stable introns, which in turn inhibit growth, forms a previously unknown node of the TOR regulatory network, which works in concert with other TORC1-dependent and TORC1-independent pathways to control growth in *S. cerevisiae* (Fig. 5g).

## Discussion

The ease by which we were able to observe accumulation of stable introns raised the question of why they had not been detected earlier, especially when considering that *S. cerevisiae* has been subjected to countless molecular analyses. The answer to this question lies with two features of our analysis. First, we examined cells in saturated culture, whereas most analyses examine cells in log phase, a stage at which all excised introns are rapidly degraded. Second, we avoided mRNA poly(A) selection, whereas many analyses perform poly(A) selection prior to RNA-seq, which depletes excised introns because they have no poly(A) tail. In addition, we used an in-house RNA-seq protocol in which RNA was fragmented and 27- to 40-nucleotide fragments were isolated for sequencing, whereas most analyses use commercial RNA-seq kits that deplete RNAs shorter than a few hundred nucleotides, the size of the excised yeast introns. Although perhaps not essential for detecting longer stable introns, this RNA-seq protocol enabled reliable and quantitative detection of stable introns regardless of their lengths. In principle, studies that used splicing-specific microarrays to assay intron retention during stress, including rapamycin and DTT treatment, might have detected stable introns^50-52^. However, those studies focus on measurements through 40 min, with no measurements taken beyond 2 h of treatment, which might not have been enough time for detection of stable introns. Our findings on prolonged TORC1 inhibition strengthen the links between TORC1 signaling and secretory stress in yeast^43-45^, and show that short and long durations of TORC1 inhibition can result in distinct effects on some downstream biological processes—including stable-intron regulation.

The discovery of a previously unknown node of the TOR regulatory network raises mechanistic questions, including that of how stable-intron accumulation might inhibit growth. Our results imply that the stoichiometry of stable introns relative to some cellular component underlies this function. One possibility is that stable introns interact with and sequester spliceosomes to reduce the splicing activity needed for ribosome biogenesis. Supporting this idea, substrate pre-mRNAs are known to compete for limited splicing machinery in *S. cerevisiae*^52^, and we found that stable introns are associated with spliceosome components (Extended Data Table 1). Moreover, in budding yeast, sequestering spliceosomes might disproportionally effect mRNAs of ribosomal protein genes (RPGs); because of their high expression levels and frequent possession of an intron, RPG mRNAs are substrates for 90% of all splicing events in log-phase *S. cerevisiae*^53^. Although this percentage presumably decreases as cells exit log phase, stable introns could function as part of a larger regulatory network that uses inefficient splicing to help keep ribosome production low during prolonged environmental and intracellular insults^53,54^. In this scenario, a large pool of stored spliceosome components would be available for release when stress subsides, allowing for rapid induction of ribosome biogenesis.

To evaluate the plausibility of this idea, we revisited our RNA-seq data from rapamycin-treated yeast to determine whether stable introns can reach the levels sufficient to have an effect on available spliceosome components. This analysis showed that in aggregate stable-intron transcripts reached 40% of the number of U5 snRNA molecules (Extended Data Fig. 8a). In principle, essential splicing proteins could be even more limiting than these abundant RNAs. Thus, stable-intron accumulation was well within a regime in which depleting 55% of the stable-intron molecules—as in our EUHSR strain—could substantially alter spliceosome availability. Furthermore, overexpressing a stable intron (*ECM33*) in the EUHSR strain grown in rapamycin resulted in more intron retention and less RPG mRNA accumulation as compared with overexpression of a non-stable intron (*ACT1*) (Extended Data Fig. 8b and c). Future characterization of the stable-intron complex will provide more detailed information on the cellular machinery that is sequestered and might also provide insight into other mechanistic questions, such as how stable introns are protected from degradation and how this protection is biochemically coupled to TORC1 inhibition.

Although our ideas regarding the mechanism of stable-intron function require additional experimental validation, one interesting aspect of this mechanism is that it operates regardless of the stable-intron sequence or genomic origin. This observation provides an example in which the history of a noncoding RNA, i.e., how the molecule is born, can be sufficient to imbue a cellular function—independent of any primary-sequence considerations. Similar principles might also apply to other noncoding RNAs, especially those that lack primary-sequence conservation.

Our results add an unexpected dimension to possible fates and functions of spliceosomal introns within eukaryotic biology. Although noteworthy examples of intron lariats and trimmed linear introns with lives post-splicing have been reported^13^, the introns described here are unique in that they persist as excised, full-length, debranched RNAs. Moreover, their stability is specifically and dramatically regulated in response to environmental changes, and they collectively act as a newly defined type of functional ncRNA. Our focus has been in the contexts of TORC1 inhibition and saturated growth, but stable introns might also be induced and have similar functions in other conditions, such as during meiosis, a time at which ribosome biogenesis and splicing competition are known to be dynamically regulated^16,52,55^. Although intron-rich eukaryotes cannot as obviously leverage global spliceosome availability to manipulate production of specific subsets of proteins, some might still use excised linear introns to perform biological functions—perhaps in environmental conditions not extensively profiled to date. At this point, we know that excised linear introns accumulate and function in at least one eukaryotic lineage, and we would be surprised if it is the only one.

## Supplementary Information

is available in the online version of the paper.

## Acknowledgements

We thank A. Amon, C. Burge, D. Pincus, D. Sabatini, P. Sharp, and members of the Bartel and Fink labs for comments and discussion, G. Li, T. Pham, and A. Symbor-Nagrabska for experimental assistance, A. Amon, R. Loewith, S. Schreiber, and V. Vyas for reagents, the Whitehead Institute Genome Technology Core for sequencing, and the Whitehead Institute Proteomics Core Facility for mass spectrometry. This work was supported by NIH grants GM035010 (G.R.F.) and GM118135 (D.P.B.). D.P.B. is an investigator of the Howard Hughes Medical Institute.

## Author Contributions

Apart from tetrad dissections performed by G.R.F., J.T.M. performed all experiments and analyses. D.P.B. supervised with help from G.R.F. All authors contributed to the design of the study and preparation of the manuscript.

## Author Information

Sequencing data will be available at the Gene Expression Omnibus. The authors declare no competing financial interests. Correspondence and requests for materials should be addressed to D.P.B.

## METHODS

### Yeast strains and genetic manipulations

*S. cerevisiae* strains used in this study are listed in Supplementary Table 1. With the exception of strains acquired for *TOR1-1* (Fig. 3c) and TORC1^∗^ (Extended Data Fig. 6) experiments, all strains were in the BY4741 background. A *TAP42* heterozygous diploid knockout strain (Horizon Discovery) was transformed with plasmids encoding either wild-type (*TAP42*) or mutant (*tap42-11*) alleles before sporulation and tetrad dissection. Transformations were performed using standard methods. Intron deletions and endogenous BP manipulations were made using a CRISPR-Cas9 system adapted for use in *S. cerevisiae*^48,49^ and were confirmed by colony PCR and Sanger sequencing of relevant genomic loci. Clones with correct deletions were counterselected for the Cas9 plasmid by plating onto 5-FOA media before their use in experiments. This process was iterated to generate strains with multiple deleted introns. Strains that underwent 5-FOA counter-selection were confirmed to not be petite by patching to YP-glycerol plates. *his3Δ1* repair was performed by transforming BY4741 with linear *HIS3* DNA, selecting for His^+^ transformants, and confirming repair of the endogenous locus by colony PCR and Sanger sequencing. Plasmids used in this study are listed in Supplementary Table 2. Oligonucleotides used for guide RNAs and repair templates are listed in Supplementary Table 3 (IDT).

### Growth conditions and harvesting

Yeast were grown at 30°C on standard synthetic complete (SC) plate and liquid media, except experiments involving TORC1^∗^ strains (Extended Data Fig. 6), which were performed according to previously described protocols^40^, and *tap42::KanMX* strains (Extended Data Fig. 5b), which were performed at 25°C due to temperature sensitivity. SC-Ura and SC-Trp were used in some experiments to maintain selection for *URA3*- and *TRP1*-expressing plasmids. SC-His was used in experiments involving 3-aminotriazole. Rapamycin (LC Laboratories; 10 mM stock in DMSO) aliquots were stored at –80°C, and diluted to 10 μM in water immediately before use. 3-aminotriazole (Sigma-Aldrich; 1 M stock in water), cycloheximide (Sigma-Aldrich; 100 mg mL^-1^ stock in DMSO), DTT (Thermo Scientific; 1 M stock in water), and tunicamycin (Millipore-Sigma; 10 mg mL^-1^ stock in DMSO) aliquots were stored at –20°C. Cultures were seeded at OD_600_ 0.2 from overnight cultures (typically OD_600_ ~6) for growth to log phase or saturation. Log-phase cultures were harvested during early log phase, typically at OD_600_ 0.5, reached 4–5 h after seeding. Unless otherwise indicated, saturated cultures were harvested 18–20 h after seeding. In acute nutrient depletion experiments, cultures were grown to mid-log phase (OD_600_ ~1), filtered, washed two times with water, and resuspended in appropriate media (SC–glucose, SC–ammonium sulfate, SC–leucine, or SC–uracil). All cultures were rapidly harvested by vacuum filtration and flash frozen in liquid nitrogen as described^57^. Frozen pellets were mechanically lysed using a Sample Prep 6870 Freezer/Mill (Spex SamplePrep; 10 cycles of 2 min on, 2 min off at setting 10). Lysate powder was aliquoted and stored at –80°C.

### Competitive growth

WT and EUHSR strains were co-cultured in a series of batch cultures, which were performed in biological replicate. The experiment started with a 50 mL coculture in a 250 mL baffled flask seeded with an equal amount of the two strains at a combined OD_600_ of 0.2. The co-culture was then allowed to grow to confluence for 3 days. At this point, it was diluted to OD_600_ of 0.2, and growth and dilution were repeated for a total of 6 cycles. 1 mL samples were taken throughout the experiment with two time points taken each cycle, one immediately before dilution and the other 16 h after dilution. Genomic DNA was harvested using the “Bust ‘n Grab” protocol^58^, and the relative abundance of WT and EUHSR was measured by PCR across the *ECM33* and *UBC4* intron-deletion loci (primers listed in Supplementary Table 3). By comparing to a standard curve of PCR results that had been templated with defined ratios of independently isolated WT and EUHSR genomic DNA (ranging from 1:4 to 4:1), the fractional content of WT cell was determined at each time point.

### RNA-Seq

Total RNA was extracted using TRI Reagent (Ambion) according to the manufacturer’s protocol. rRNA was depleted from 5 μg of total RNA using the Ribo-Zero Gold Yeast rRNA Removal Kit (Illumina) according to the manufacturer’s protocol. RNA-seq libraries were prepared as described^59^ and sequenced on the Illumina HiSeq platform using 40 bp single reads. A detailed protocol is available at http://bartellab.wi.mit.edu/protocols.html.

### RNA-Seq Analyses

Reads were trimmed of adaptor sequence using cutadapt^60^. Trimmed reads were aligned to the *S. cerevisiae* genome (R64-1-1, downloaded from www.yeastgenome.org) using STAR^61^ (v2.4) with the parameters “‐‐alignIntronMax 1000 ‐‐sjdbOverhang 31 ‐‐outSAMtype BAM SortedByCoordinate ‐‐quantMode GeneCounts” and with “‐‐sjdbGTFfile” supplied with transcript annotations. Intron annotations were constructed from the UCSC Genome browser and the Ares lab yeast intron database (http://intron.ucsc.edu/yeast4.3/), with the Ares lab database^14,15^ providing branch point annotations used in subsequent analyses. RNA-seq visualization was performed using IGV (v.2.3.57)^62,63^.

Untemplated nucleotides were found and cataloged by extracting reads from BAM files that uniquely mapped to the sense strand of an intron but which also contained soft-clipped, non-mapping nucleotides at the 3′ end of the read. The Mixture-of-Isoforms^31^ (MISO, v.0.5.4) framework was used to quantify relative expression of three potential isoforms for every intron: spliced (intron degraded), spliced (intron stable), and intron retained. MISO requires uniform read length as input. As such, 3′ adaptor sequence was removed by trimming a constant 8 nt from every read. Based on the distribution of prior cutadapt-mediated trimming, this removed all 3′ adaptor sequence from >98% of reads. Retained intron GFF events were constructed using “gff_make_annotation” from rnaseqlib (http://rnaseqlib.readthedocs.io/en/clip/). Stable intron GFF events were made by directly modifying the retained intron GFF events to instead include “intron only” as a potential outcome of splicing. Although reads that supported a stable-intron splicing event necessarily could also support a retained-intron splicing event, in practice, the greater abundance of most stable introns relative to flanking exons enabled MISO to identify stable intron as the predominant isoform for many introns in stable-intron–inducing conditions.

To be identified as a stable intron expressed in a given condition, accumulation of the intron had to exceed thresholds calibrated on experimentally validated cases. First, intron accumulation (transcripts per million, TPM) in the stable-intron–inducing condition had to be greater than 50% of exon accumulation (TPM). Second, the intron accumulation (TPM) in the stable-intron–inducing condition had to be more than twice that of its accumulation (TPM) in log phase (assigning a pseudocount of 0.1 reads to introns with 0 reads). Third, the ratio of intron:exon accumulation in the stable-intron–inducing condition had to be greater than 4-fold that of the intron:exon ratio in log phase. Imposing these thresholds eliminated many false-positives, including those attributed to intronic snoRNA expression or constitutively poor splicing. For introns exceeding these accumulation thresholds, at least one of the following two additional criteria were required for annotation as a stable intron: 1) ≥ 2 terminal adenylylated reads mapping to the 3′ terminus of the intron, or 2) MISO-based support for preferential accumulation of the stable-intron isoform in the stable-intron–inducing condition (Bayes factor > 30 when compared to log phase). These criteria identified conservative sets of stable introns expressed in a given condition, erring towards reducing false-positive identifications.

A search for motifs enriched in stable introns was performed using the MEME Suite^64^ with remaining introns as the background set. *k*-mer frequencies were generated with fasta-get-markov program from the MEME Suite. In addition, a search for enrichment of position-specific motifs was performed using *k*pLogo^65^ with remaining introns as the background set.

### Growth curves

Growth curves were collected using Nunc Edge 2.0 96-well plates in a Multiskan GO Microplate Spectrophotometer. Wells were seeded at OD_600_ = 0.2 from overnight cultures, and plates were incubated at 30 °C. Absorbance was read every 5 min with shaking on the “background” setting, cycling between 1 min on and 1 min off. Every strain tested on each plate was run in technical triplicate. Single wells were censored if artefactual spikes in OD_600_ attributable to bubbles or condensation were observed. Strains were censored if more than one well was censored. Biological replicates for each strain included at least two independently derived transformants. A replicate of parental BY4741 was included on every plate as a control strain, and SC media was included on every plate as a control condition for every strain being assayed. Growth rate was calculated from the log-linear portion of each growth curve. Growth curves were analyzed with SkanIt (v3.2).

### RNA blots

10 μg of total RNA for each sample was resolved on a denaturing polyacrylamide gel and transferred onto a Hybond membrane (GE Healthcare) using a semi-dry transfer cell. Because UV crosslinking is biased against shorter RNAs, EDC (*N*-(3-dimethylaminopropyl)-*N*′-ethylcarbodiimide; Sigma-Aldrich) was used to chemically crosslink 5′ phosphates to the membrane^66^. Blots were hybridized to radio-labeled DNA probes. Probe oligonucleotides are listed in Supplementary Table 4. More details of this protocol are available at http://bartellab.wi.mit.edu/protocols.html. For experiments in Fig. 3a, RNAs were instead resolved on a glyoxal agarose gel and transferred overnight onto a Nytran SuPerCharge Turboblotter membrane (GE Healthcare). RNA blot data were analyzed with ImageQuant TL (v8.1.0.0).

### Sedimentation velocity

Crude lysates were prepared by re-suspending an aliquot of thawed lysate powder (500–800 μL of loosely packed powder) in 1 mL of lysis buffer (10 mM Tris-HCl [pH 7.4], 5 mM MgCl_2_, 100 mM KCl, 1% Triton X-100, 1% Sodium Deoxycholate, 2 mM DTT, 20 U/ml SUPERase•In [Ambion], cOmplete EDTA-free Protease Inhibitor Cocktail [Roche]). The lysates were placed on a rotator mixer at 4 °C for 5 min to allow for re-suspension. Following brief vortexing, lysates were centrifuged at 1,300 × g for 10 min, and the supernatant loaded onto a 12.5 mL linear 10–30% (w/v) sucrose gradient (20 mM HEPES-KOH [pH 7.4], 5 mM MgCl_2_, 100 mM KCl, 2 mM DTT, 20 U/ml SUPERase•In). Gradients were centrifuged in a pre-chilled SW-41 Ti rotor at 38,000 rpm for 4 hr at 4 °C. Gradients were fractionated using the Piston Gradient Fractionator (Biocomp) in 1 mL fractions. For RNA analysis, total RNA was extracted from a portion of each fraction using TRI Reagent. RNA blots were performed as described above, except RNA loading was normalized by percent of gradient fraction rather than by RNA concentration. For pull-downs, fractions were flash frozen and stored at −80°C.

### Ectopic intron expression, affinity purification, and mass spectrometry

Intron overexpression constructs were based on SC-Ura-selectable pRSII416 (CEN) or pRSII426 (2μ) vectors^67^ (Extended Data Fig. 3c, Supplementary Table 2). Introns were inserted 49 nucleotides downstream of the *URA3* start codon by Gibson assembly^68^. Proper splicing was confirmed by RNA blot and cell viability, as cells unable to produce Ura3p through intron removal could not grow on SC-Ura media.

Pull-down experiments were performed in the *ECM33Δintron* strain. Introns were purified utilizing two MS2 hairpins (2×MS2) inserted in various positions within the *ECM33* intron (Supplementary Table 5). To minimize potential effects on splicing, sequence within 60 nt of the 5′ splice site and 80 nt of the branch point was kept constant across all constructs. The 2×MS2 hairpin sequence was based on CRISPR RNA scaffold designs^69^. 3×FLAG-tagged MS2 coat protein with a C-terminal nuclear localization signal (FLAG-MCP) was co-expressed from the same construct as MS2-tagged introns.

We were unable to purify intact introns from supernatant of saturated cultures due to increased endogenous RNase activity in saturated cultures^70^. To circumvent this, sucrose gradient fractions containing the intron of interest (typically fractions 6–9, determined by RNA blot) were pooled and used as the starting material for purification. ANTI-FLAG M2 Magnetic Beads (Sigma-Aldrich) (20 μL of packed bead volume) were equilibrated and washed twice in 10 volumes of buffer 1 (100 mM KCl, 20 mM HEPES KOH [pH 7.9], 1% Triton X-100, 20 U/ml SUPERaseI•n, and cOmplete EDTA-free Protease Inhibitor Cocktail). 400 μL of the pooled fractions were added to the beads and incubated on a rotator mixer for 30 min at 4 °C. Remaining at 4 °C, the beads were washed twice in 10 volumes of buffer 1 and twice in 10 volumes of buffer 2 (200 mM KCl, 20 mM HEPES KOH [pH 7.9], 1% Triton X-100, 20 U/ml SUPERase•In, and cOmplete EDTA-free Protease Inhibitor Cocktail). Bound FLAG-MCP was eluted from beads with 30 μL of 150 ng/μL 3× FLAG peptide (Sigma-Aldrich) on a rotator mixer for 30 min at 4 °C. The eluents were precipitated with TCA, digested with trypsin, and labeled with TMTs to allow for quantitative comparisons between 6 total samples (3 control and 3 test samples). Peptides were analyzed by liquid chromatography-tandem mass spectrometry (LC-MS/MS) using an Orbitrap Elite (Thermo Fisher) coupled with a NanoAcquity UPLC system (Waters). Peptides were identified using SEQUEST and data analyzed using PEAKS Studio (Bioinformatics Solutions).

### Experimental design and reproducibility

No statistical methods were used to predetermine sample size. Growth curve cultures were randomized by permutation of strain placement on 96-well plates across experiments. Edge effects were not significant within the time measurements were taken. The investigators were not blinded to allocation during experiments and outcome assessment. RNA-seq results for biological replicates correlated well (log-phase culture, *R*^2^ = 0.98 [mRNA, n = 5898]; saturated culture, *R*^2^ = 0.90 [mRNA, n = 6211] and 0.78 [stable introns, n = 29]).

## Extended Data Figures

**Extended Data Figure 1.**
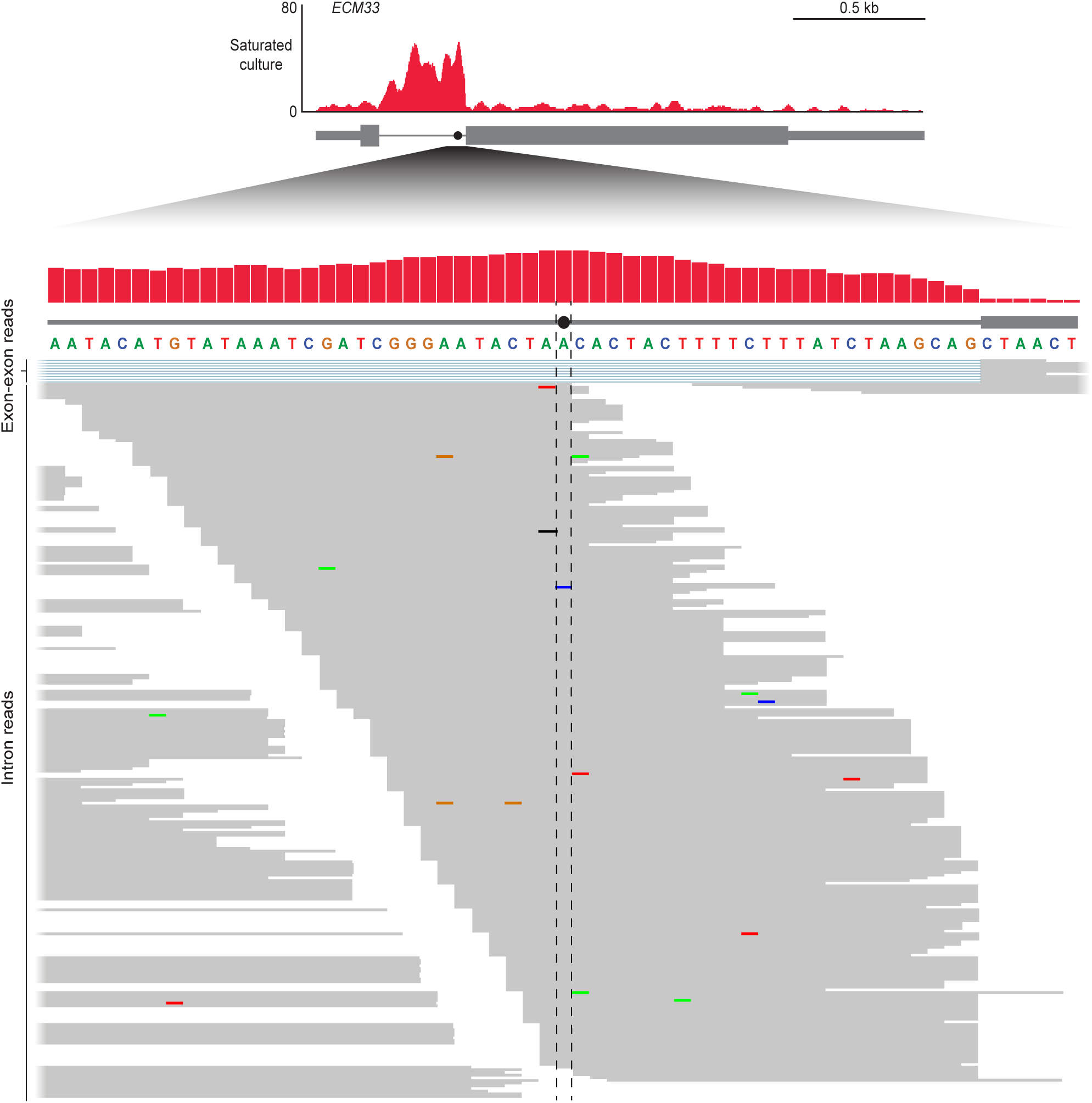
RNA-seq coverage across the BP and to the 3′SS of the *ECM33* intron. Shown is the pileup of reads mapping to the 61-nucleotide region centered on the *ECM33* BP, replotted from Fig. 1b (red, top). The position of the BP (closed circle with flanking dashed lines) is indicated on the intron (thick gray line) and relative to 3′ exon (gray box). Below, all reads mapping uniquely to this region are shown (thin gray lines). Reads mapping across the exon-exon junction are colored blue in the region of the excised intron and are shown above the other reads. Mismatched nucleotides within reads are indicated with colored bars, with color of the bar indicating the identity of the mismatch. Terminal untemplated nucleotides have been clipped from reads.

**Extended Data Figure 2.**
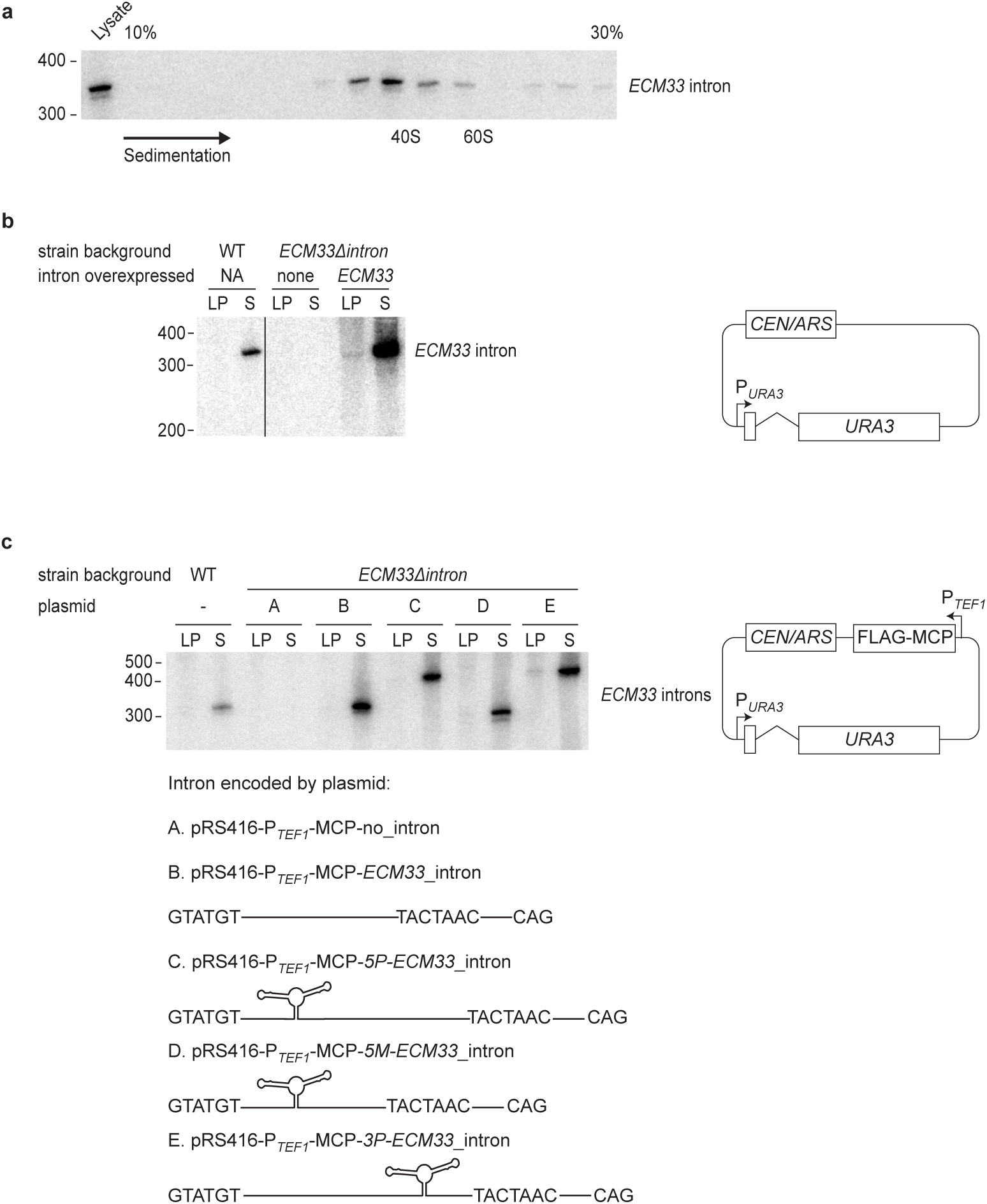
Stable-intron sedimentation and expression constructs for pull-down and mass spectrometry. **a**, Co-sedimentation of the *ECM33* stable intron with large ribonucleoprotein complexes. Cleared lysate from a saturated yeast culture was fractionated by sedimentation into a 10-30% glucose gradient, and RNA was extracted from each fraction. Shown is an RNA blot that resolved 25% of the RNA from each fraction and was probed for the *ECM33* intron. Fractions are oriented with increasing sedimentation left-to-right. RNA was also extracted from 12.5% of the total lysate before sedimentation, of which 50% was loaded for comparison (left lane). Sedimentation of 40S and 60S complexes are marked based on sedimentation of the ribosome subunits. Migration of markers with lengths indicated (nucleotides) is shown at the left. **b**, Endogenous behavior of stable introns ectopically expressed from the *URA3* gene. Shown is an RNA blot probed for the *ECM33* intron after resolving total RNA from cultures expressing the *ECM33* intron from the endogenous *ECM33* locus (WT, left lanes), cultures from a strain in which the intron had been deleted (*ECM33Δintron*, middle lanes), and cultures of this deletion strain ectopically expressing the *ECM33* intron spliced from the plasmid-borne *URA3* reporter gene (right lanes, pRS416-ECM33_intron plasmid shown on right, P_*URA3*_, *URA3* promoter; *CEN/ARS*, low-copy origin of replication). Interior lanes not relevant to this experiment were removed from this image (vertical line). Otherwise, this panel is as in Fig. 1c. **c**, Endogenous behavior of stable introns with MS2 hairpins inserted to be used as affinity tags for pull-downs. Five different plasmids with a common backbone (right; P_*TEFI*_, *TEF1* promoter; *FLAG-MCP*, coding region of FLAG-tagged MS2 coat protein) each expressed *URA3* with a different variant of the *ECM33* intron (variants A–E, schematized below). These plasmids were each expressed in the strain that lacked an endogenous *ECM33* intron (*ECM33Δintron*). The RNA blot resolved total RNA from the indicated cultures and was probed for a sequence common to the intron variants; otherwise as in Fig. 1c. The 2×MS2 hairpin region is 90 nucleotides long, and the expected linear-intron sizes were: A, no intron; B, 330 nucleotides; C, 420 nucleotides; D, 300 nucleotides; E, 420 nucleotides.

**Extended Data Figure 3.**
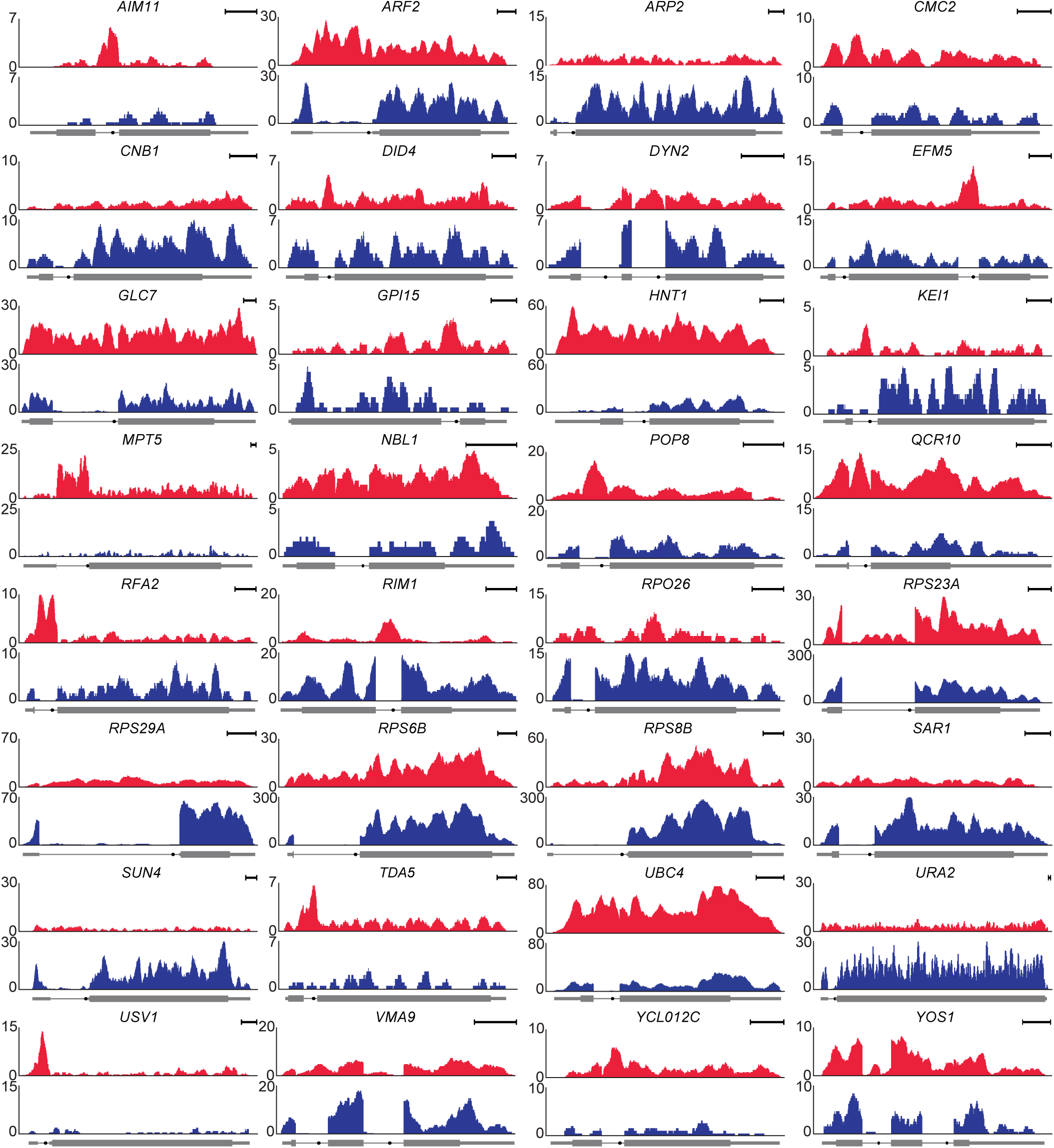
RNA-seq profiles of genes with stable introns. Profiles from the stable-intron-inducing condition (red) and log-phase culture (blue) are shown for the indicated 32 genes with confidently identified stable introns not already depicted in Figure 1a. For all but four of these, the profile of the stable-intron-inducing condition is from RNA-seq of the saturated-liquid sample. The four exceptions were not confidently identified in saturated liquid (Extended Data Table 2); for these, the profile of the stable-intron–inducing condition is from either the rapamycin-treated sample (*DYN2_2* and *TDA5*) or the saturated lawn (*RPO26* and *RPS8B*). Scale bars, 100 nucleotides. Otherwise, this panel is as in Fig. 1a.

**Extended Data Figure 4.**
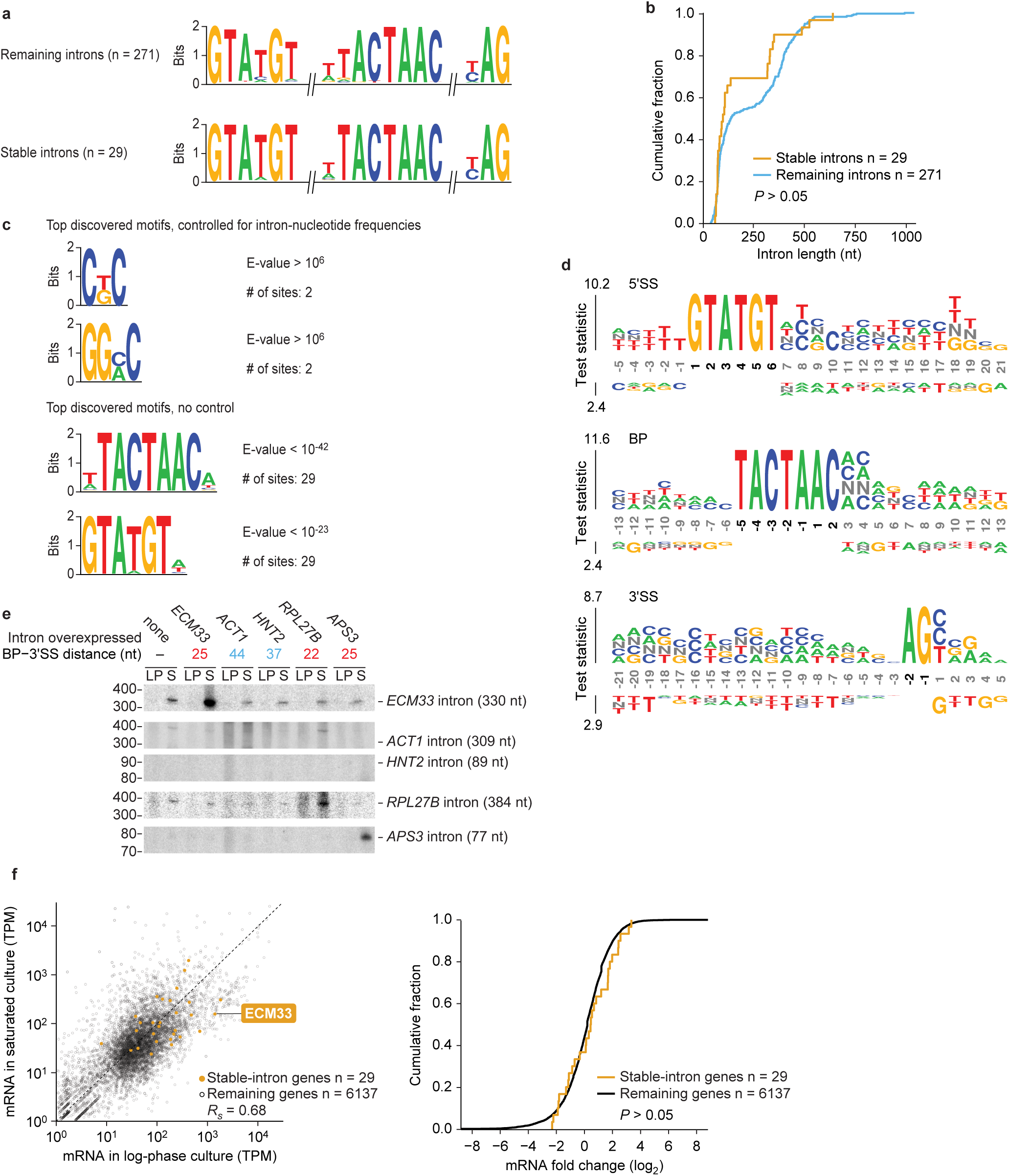
Stable introns are indistinguishable from other introns in many respects. **a**, Similar splicing motifs compared to other introns. Plotted are information-content logos of splicing motifs (6-mer 5′SS, 8-mer BP, and 3-mer 3′SS) for stable introns (bottom) and other introns (top). **b**, Similar length distribution compared to other introns. Plotted are cumulative distributions of intron lengths (*P* > 0.05, one-tailed Kolmogorov–Smirnov test). **c**, No significantly enriched motifs within stable introns. Plotted are stable-intron sequence motifs discovered by MEME^64^ either controlling for nonstable intron *k*-mer frequencies (top) or without controlling (bottom). No significant motifs were discovered in stable introns when *k* ≥ 6. The motifs discovered without controlling for *k*-mer frequencies matched the canonical BP and 5′SS motifs (see **a**) BP and 5′SS motifs were also the only significantly enriched motifs discovered when k ≤ 5. **d**, No enriched positional *k*-mer motifs detected by *k*pLogo^65^ in stable introns. Plotted are the most enriched *k*-mers at positions relative to 5′SS (top), BP (middle), and 3′SS (bottom) comparing stable introns to unstable introns. Stacked nucleotides at a position represent the most significant motif starting at that position. The height is scaled relative to the significance of the motif, as determined by the one-sided binomial test statistic (y axes). Black numbers indicate invariant nucleotides occurring > 95% of the time at the position. No *k*-mers were significantly enriched when using a Bonferroni-corrected *P* of 0.01. **e**, Support for a role of BP–3′SS distance in specifying stable-intron formation. The indicated introns were ectopically expressed from the *URA3* splicing construct (Extended Data Fig. 2b). Shown are results from an RNA blot that resolved total RNA from cultures overexpressing the indicated introns and was probed for the indicated introns (length of excised intron in parentheses). The *ECM33* and *RPL27B* introns were probed sequentially, and then the *ACT1, HNT2,* and *APS3* introns were probed concurrently. *ACT1* probe was validated on synthetic transcripts resembling the *ACT1* intron, which were produced by *in vitro* transcription (not shown). Migration of markers with lengths indicated (nucleotides) is at the left. **f**, Expression of mRNAs from genes containing stable introns. Left, relationship between the expression in log-phase and saturated cultures (expression cutoff, 1 transcript per million [TPM]). Points for genes expressing stable introns are indicated (orange), and the one for *ECM33* is labeled. Right, comparison of the expression results of genes with stable introns to those of the remaining genes. Plotted are the cumulative distributions of log_2_-fold-changes in expression observed between log-phase and saturated cultures, which shows no significant difference between stable-intron genes and other genes. (*P* > 0.05, two-tailed Kolmogorov–Smirnov test).

**Extended Data Figure 5.**
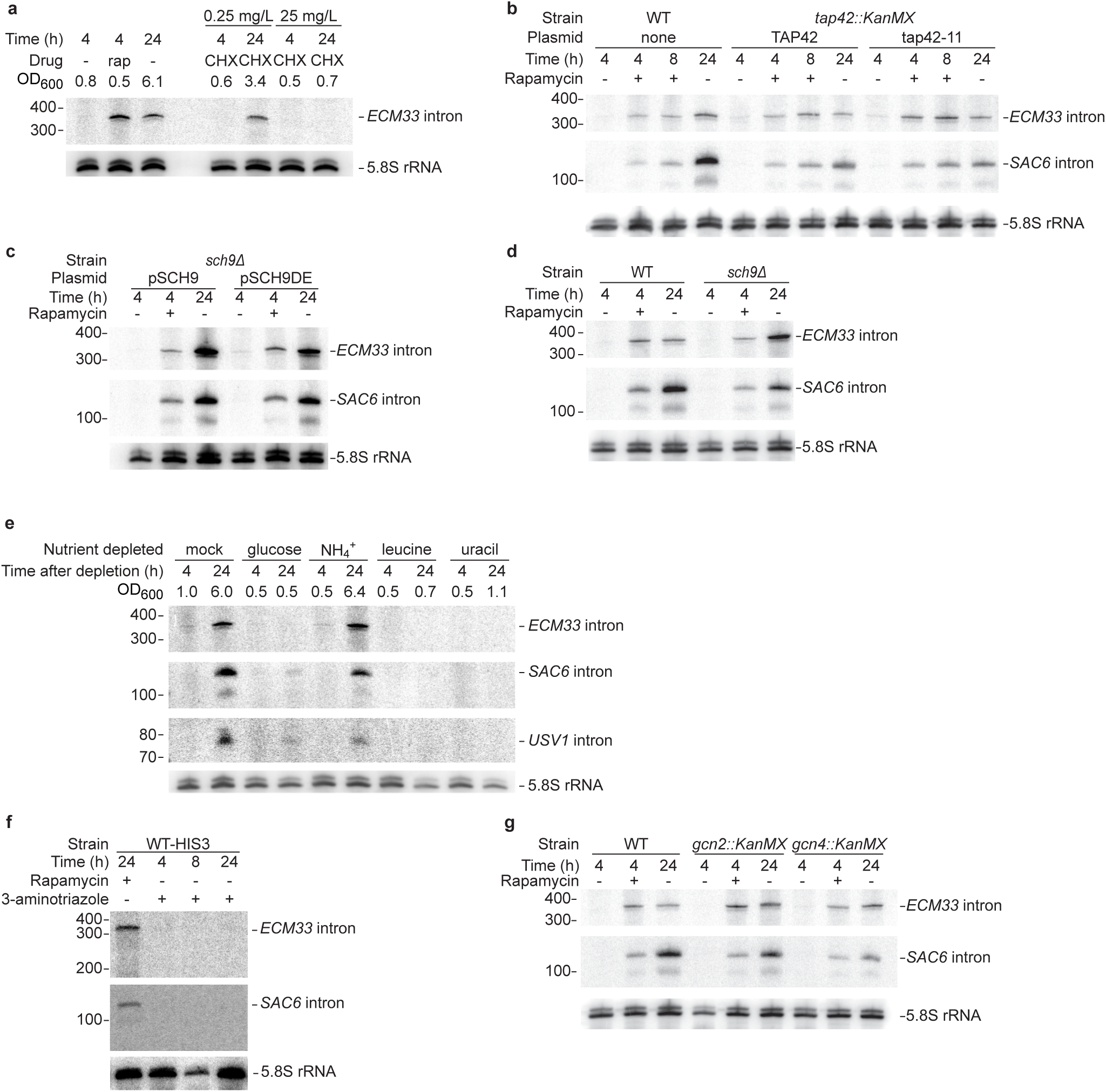
Assessing aspects of TORC1 regulation on stable-intron expression. **a**, Inability of TORC1-independent inhibition of protein synthesis to prematurely induce stable-intron accumulation. The left lanes show a replicate of Fig. 3b, and the right lanes show results after treatment with either low (0.25 mg L^-1^) or high (25 mg L^-1^) concentrations of cycloheximide. As indicated by OD_600_ at harvest, the mild cycloheximide treatment allowed the culture to reach an OD600 of 3.4 after 24 h, which was equivalent to the OD_600_ of 10 h without cycloheximide (Fig. 3a). **b**, Dispensibility of TORC1-responsive Tap42 for stable-intron formation. Samples were grown at 25°C due to temperature sensitivity of the *tap42-11* allele. Additionally, due to slower growth, the duration of pre-growth for −/+ rapamycin data points was extended to 5.5 h, such that the change in OD_600_ of the pre-growth sample matched that of WT cultures grown at 30°C for 4 h. Otherwise, this panel is as in Fig. 3b. **c**, Dispensibility of TORC1-responsive Sch9 for stable-intron formation. Samples were grown in SC-Trp media to maintain plasmids that either rescued (*SCH9*) or did not rescue (*SCH9DE*) Sch9 activity. Because of the slower growth of the *sch9Δ pRS414::SCH9* and *sch9Δ pRS414::SCH9DE* strains, possibly due to the requisite–Ura media, the duration of pre-growth for −/+ rapamycin data points was extended to 5 h, such that the change in OD_600_ of the pre-growth sample matched that of WT cultures grown for 4 h. Otherwise, this panel is as in Fig. 3b. **d**, Dispensibility of Sch9 for stable-introns formation. The left lanes show a replicate of Fig. 3b, and the right lanes show the same experiments performed in the *sch9Δ* strain. **e**, Inability of acute deprivation of select nutrients to induce accumulation of stable introns. To prevent contamination of starting cultures with stable introns contributed by inoculum, cultures were seeded at OD_600_ = 0.2 from an overnight culture that was allowed to grow to mid-log phase, collected by vacuum filtration, washed in water, and resuspended in fresh media lacking the indicated nutrients. Cultures were harvested after the indicated times and analyzed as in Fig. 1c, except the RNA blot was sequentially reprobed for the *ECM33, SAC6*, and *USV1* introns. As indicated by OD_600_ at harvest, the sample deprived of ammonium sulfate, the main nitrogen source, was still able to reach a high density after 24 h. **f**, Stable-intron accumulation despite inhibition of the GAAC pathway. Mid-log cultures were treated with either 100 nM rapamycin or 50 mM 3-aminotriazole for the indicated times. Samples were grown in SC-His to force inhibition of histidine biosynthesis by 3-aminotriazole. **g**, Stable-intron accumulation despite inability to induce the GAAC pathway. The left lanes show a replicate of Fig. 3b, and the middle and right lanes show the same experiments performed in the *gcn2Δ* and *gcn4Δ* strains.

**Extended Data Figure 6.**
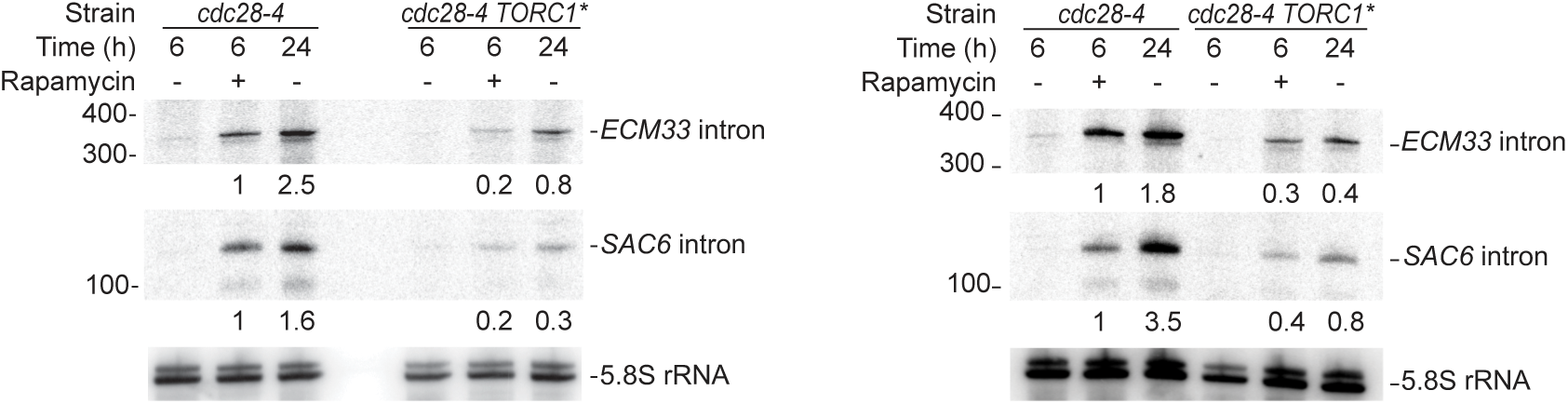
The influence of hyperactive TORC1 on stable-intron formation. The left and right panels are biological replicates comparing *ECM33* nd *SAC6* intron levels in a strain with hyperactive TORC1 (TORC1^∗^) to those in a strain-background control (*cdc28-4*). Growth conditions were was described for these strains^40^. Overexpression of Sfp1 was induced 1 h prior to −/+ rapamycin treatment. Due to the decreased growth rate of these strains in the requisite conditions, −/+ rapamycin samples were harvested after 6 h of treatment rather than after 4 h of treatment. Numbers below *ECM33* and *SAC6* intron blots indicate the level of each intron normalized to the level in the *cdc28-4* +rapamycin samples, after first normalizing all lanes to the 5.8S rRNA loading control (set to 1).

**Extended Data Figure 7.**
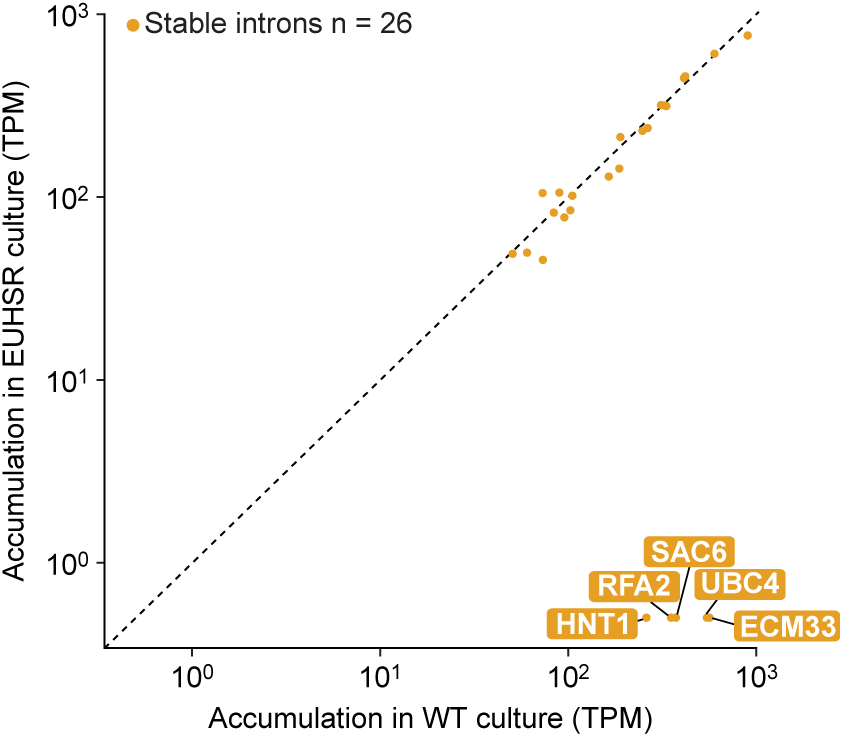
Assessing stable-intron expression in a EUHSR culture. No evidence for compensatory stable-intron expression after genomic deletion of five stable introns. The scatter plot shows the relationship between intron accumulation (TPM, transcripts per million) in rapamycin-treated WT culture and in rapamycin-treated EUHSR culture. The dashed line is placed at *x = y*. Stable introns are indicated (closed orange circles). Points for introns deleted from the EUHSR genome (*ECM33, UBC4, HNT1, SAC6*, and *RFA2*) are labeled. All introns were pseudo-counted at 0.5 TPM.

**Extended Data Figure 8.**
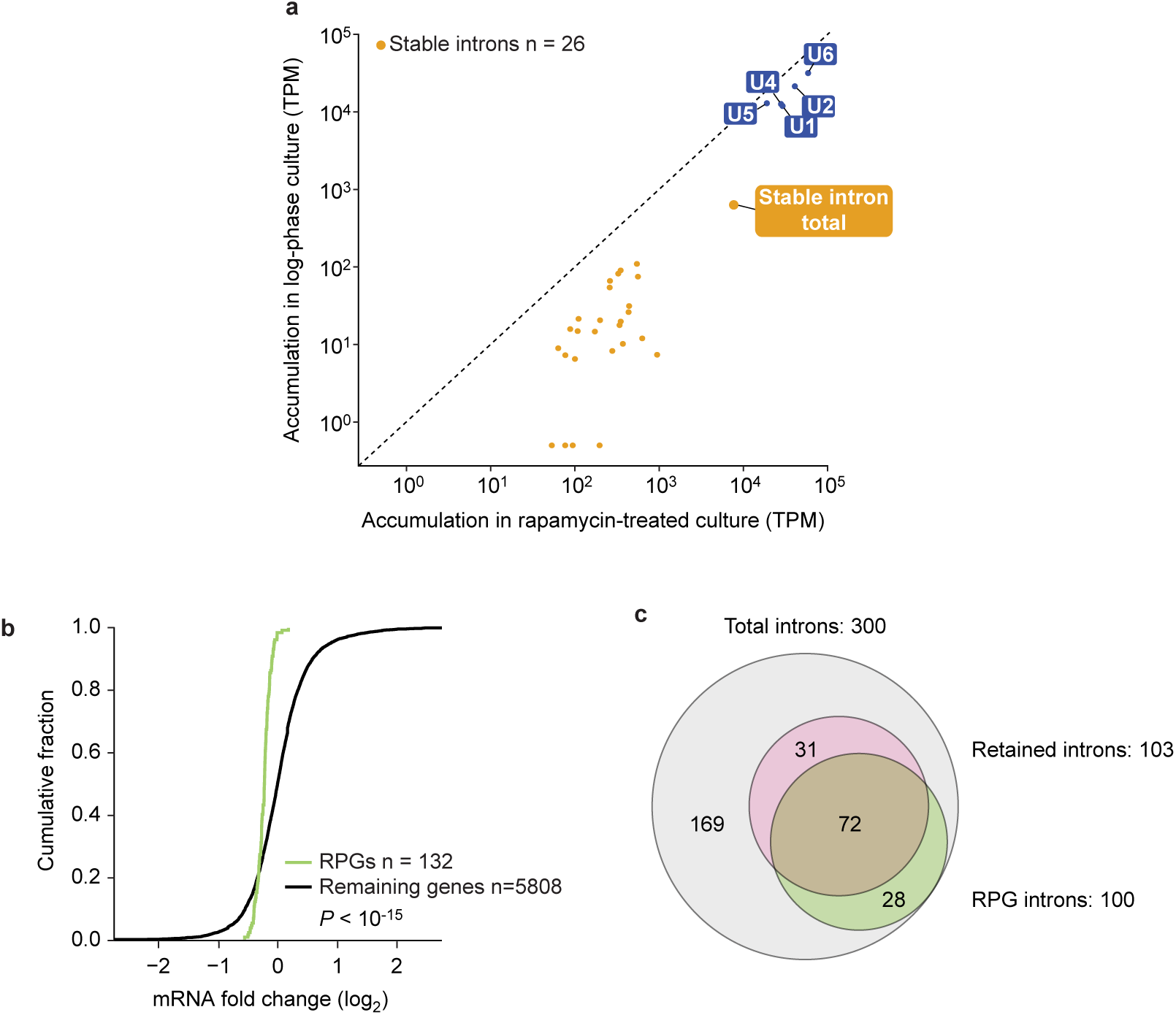
Evidence for spliceosome sequestration and control of ribosome production by stable introns. **a**, Aggregate stable-intron accumulation approaching that of spliceosomal RNAs. Plotted are levels of stable introns and spliceosomal RNAs (labeled closed blue circles), comparing levels in log-phase WT culture to those in rapamycin-treated WT culture. Also plotted is the aggregate stable-intron abundance (closed orange circle, “stable intron total”). Otherwise, as in Extended Data Fig. 7. **b**, Reduced RPG mRNA expression when overexpressing a stable intron in rapamycin-treated EUHSR culture. Plotted are the cumulative distributions of log2-fold-changes in mRNA expression observed between a EUHSR culture with stable-intron (*ECM33*) ectopic expression and a EUHSR culture with control-intron (*ACT1*) ectopic expression. The distribution of changes for mRNAs of RPGs (green) differed from that other genes (black), with generally lower expression of RPGs in the culture overexpressing the stable intron (*P* < 10^-15^, one-tailed Kolmogorov-Smirnov test; expression cutoff, 1 TPM in both samples). **c**, Less efficient splicing, as detected by increased intron retention, when overexpressing a stable intron in rapamycin-treated EUHSR culture. When analyzing data set of **b**, 103 genes had significantly more intron retention when ectopically expressing the stable intron compared to when expressing the control intron (MISO, Bayes factor > 100). The Venn diagram shows the overlap between these genes with increased intron retention and intron-containing RPGs (*P* < 10^-21^, hypergeometric test).

**Extended Data Table 1.** Proteins associated with stable introns. Shown are results for the ten proteins consistently enriched ≥ 2 fold in stable-intron pull-down eluates. Three control samples without a tagged intron and three test samples with a unique tagged introns (Extended Data Fig. 2, Supplementary Table 5) were simultaneously analyzed by quantitative mass spectrometry. These ten proteins were the only proteins enriched ≥ 2fold in each of the nine possible pairwise comparisons between test and control samples. The identities of these proteins were consistent with the excised and debranched introns being part of a complex resembling the ILS complex, in that all ten are known components of the ILS identified through biochemical studies^29^, and most (all but Brr2, Hsh49, Prp9, and Rse1) have also been identified in a cryo-EM structure of the ILS complex ^71^. Moreover, several early spliceosome components (Luc7, Prp3, Snp1, and Snu13) as well as an essential disassembly factor (Prp43) were identified across all samples but not enriched in tagged-intron eluates. Enrichment values were those reported by PEAKS Studio, which are reported relative to values of control 2.

| Protein name | Enrichment values |  |  |  |  |  |  |  |  |
| --- | --- | --- | --- | --- | --- | --- | --- | --- | --- |
|  | Control 1 | Control 2 | Control 3 | Test 1 | Test 2 | Test 3 | Mean control | Mean test | Fold difference |
| Prp9 | 0.00 | 1.00 | 0.57 | 4.84 | 6.34 | 4.83 | 0.52 | 5.34 | 10.20 |
| Hsh49 | 0.60 | 1.00 | 1.20 | 5.67 | 5.15 | 5.24 | 0.93 | 5.35 | 5.74 |
| Cef1 | 0.93 | 1.00 | 0.99 | 3.45 | 6.04 | 5.79 | 0.97 | 5.09 | 5.23 |
| Rse1 | 0.66 | 1.00 | 0.90 | 3.17 | 4.16 | 5.80 | 0.85 | 4.38 | 5.13 |
| Bud31 | 0.68 | 1.00 | 0.96 | 3.96 | 3.96 | 4.16 | 0.88 | 4.03 | 4.58 |
| Prp45 | 0.88 | 1.00 | 0.71 | 3.12 | 3.77 | 3.86 | 0.86 | 3.58 | 4.15 |
| Prp19 | 0.87 | 1.00 | 1.05 | 3.28 | 4.03 | 4.20 | 0.97 | 3.84 | 3.94 |
| Brr2 | 0.78 | 1.00 | 0.69 | 2.22 | 3.21 | 3.66 | 0.82 | 3.03 | 3.68 |
| Prp8 | 0.75 | 1.00 | 0.73 | 2.00 | 2.63 | 3.46 | 0.83 | 2.70 | 3.26 |
| Smb1 | 0.67 | 1.00 | 1.10 | 2.49 | 2.80 | 2.69 | 0.92 | 2.66 | 2.88 |

**Extended Data Table 2.**
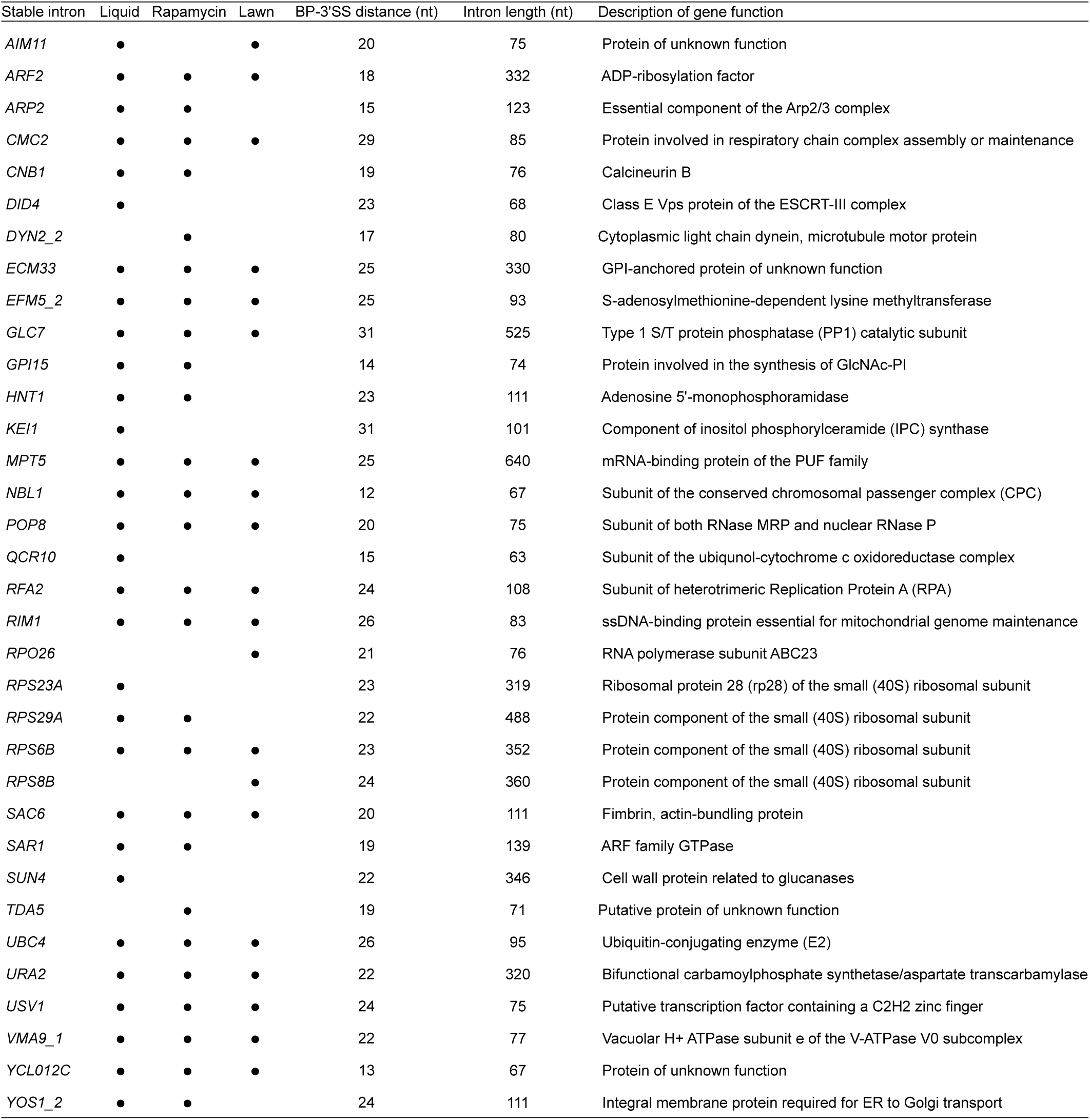
Stable introns identified in saturated cultures. Description of gene function is from YeastMine (https://yeastmine.yeastgenome.org/). Liquid, saturated-liquid sample; Lawn, saturated-lawn sample; Rapamycin, rapamycin-treated sample.

| Stable intron | Liquid | Rapamycin | Lawn | BP-3'SS distance (nt) | Intron length (nt) | Description of gene function |
| --- | --- | --- | --- | --- | --- | --- |
| *AIM11* | • |  | • | 20 | 75 | Protein of unknown function |
| *ARF2* | • | • | • | 18 | 332 | ADP-ribosylation factor |
| *ARP2* | • | • |  | 15 | 123 | Essential component of the Arp2/3 complex |
| *CMC2* | • | • | • | 29 | 85 | Protein involved in respiratory chain complex assembly or maintenance |
| *CNB1* | • | • |  | 19 | 76 | Calcineurin B |
| *DID4* | • |  |  | 23 | 68 | Class E Vps protein of the ESCRT-III complex |
| *DYN2_2* |  | • |  | 17 | 80 | Cytoplasmic light chain dynein, microtubule motor protein |
| *ECM33* | • | • | • | 25 | 330 | GPI-anchored protein of unknown function |
| *EFM5_2* | • | • | • | 25 | 93 | S-adenosylmethionine-dependent lysine methyltransferase |
| *GLC7* | • | • | • | 31 | 525 | Type 1 S/T protein phosphatase (PP1) catalytic subunit |
| *GPI15* | • | • |  | 14 | 74 | Protein involved in the synthesis of GlcNAc-PI |
| *HNT1* | • | • |  | 23 | 111 | Adenosine 5'-monophosphoramidase |
| *KEI1* | • |  |  | 31 | 101 | Component of inositol phosphorylceramide (IPC) synthase |
| *MPT5* | • | • | • | 25 | 640 | mRNA-binding protein of the PUF family |
| *NBL1* | • | • | • | 12 | 67 | Subunit of the conserved chromosomal passenger complex (CPC) |
| *POP8* | • | • | • | 20 | 75 | Subunit of both RNase MRP and nuclear RNase P |
| *QCR10* | • |  |  | 15 | 63 | Subunit of the ubiquinol-cytochrome c oxidoreductase complex |
| *RFA2* | • | • | • | 24 | 108 | Subunit of heterotrimeric Replication Protein A (RPA) |
| *RIM1* | • | • | • | 26 | 83 | ssDNA-binding protein essential for mitochondrial genome maintenance |
| *RPO26* |  |  | • | 21 | 76 | RNA polymerase subunit ABC23 |
| *RPS23A* | • |  |  | 23 | 319 | Ribosomal protein 28 (rp28) of the small (40S) ribosomal subunit |
| *RPS29A* | • | • |  | 22 | 488 | Protein component of the small (40S) ribosomal subunit |
| *RPS6B* | • | • | • | 23 | 352 | Protein component of the small (40S) ribosomal subunit |
| *RPS8B* |  |  | • | 24 | 360 | Protein component of the small (40S) ribosomal subunit |
| *SAC6* | • | • | • | 20 | 111 | Fimbrin, actin-bundling protein |
| *SAR1* | • | • |  | 19 | 139 | ARF family GTPase |
| *SUN4* | • |  |  | 22 | 346 | Cell wall protein related to glucanases |
| *TDA5* |  | • |  | 19 | 71 | Putative protein of unknown function |
| *UBC4* | • | • | • | 26 | 95 | Ubiquitin-conjugating enzyme (E2) |
| *URA2* | • | • | • | 22 | 320 | Bifunctional carbamoylphosphate synthetase/aspartate transcarbamylase |
| *USV1* | • | • | • | 24 | 75 | Putative transcription factor containing a C2H2 zinc finger |
| *VMA9_1* | • | • | • | 22 | 77 | Vacuolar H<sup>+</sup> ATPase subunit e of the V-ATPase V0 subcomplex |
| *YCL012C* | • | • | • | 13 | 67 | Protein of unknown function |
| *YOS1_2* | • | • |  | 24 | 111 | Integral membrane protein required for ER to Golgi transport |

## References

1 Koonin, E. V. The origin of introns and their role in eukaryogenesis: a compromise solution to the introns-early versus introns-late debate? Biol Direct 1, 22, doi:10.1186/1745-6150-1-22 (2006).

2 Irimia, M. & Roy, S. W. Origin of spliceosomal introns and alternative splicing. Cold Spring Harbor perspectives in biology 6, doi:10.1101/cshperspect.a016071 (2014).

3 Grabowski, P. J., Padgett, R. A. & Sharp, P. A. Messenger RNA splicing in vitro: an excised intervening sequence and a potential intermediate. Cell 37, 415-427 (1984).

4 Padgett, R. A., Konarska, M. M., Grabowski, P. J., Hardy, S. F. & Sharp, P. A. Lariat RNA’s as intermediates and products in the splicing of messenger RNA precursors. Science 225, 898-904 (1984).

5 Ruskin, B., Krainer, A. R., Maniatis, T. & Green, M. R. Excision of an intact intron as a novel lariat structure during pre-mRNA splicing in vitro. Cell 38, 317-331 (1984).

6 Rodriguez, J. R., Pikielny, C. W. & Rosbash, M. In vivo characterization of yeast mRNA processing intermediates. Cell 39, 603-610 (1984).

7 Domdey, H. et al. Lariat structures are in vivo intermediates in yeast pre-mRNA splicing. Cell 39, 611-621 (1984).

8 Ruskin, B. & Green, M. R. Specific and stable intron-factor interactions are established early during in vitro pre-mRNA splicing. Cell 43, 131-142 (1985).

9 Sharp, P. A. et al. in Cold Spring Harbor symposia on quantitative biology. 277-285 (Cold Spring Harbor Laboratory Press).

10 Chapman, K. B. & Boeke, J. D. Isolation and characterization of the gene encoding yeast debranching enzyme. Cell 65, 483-492 (1991).

11 Arenas, J. & Hurwitz, J. Purification of a RNA debranching activity from HeLa cells. Journal of Biological Chemistry 262, 4274-4279 (1987).

12 Black, D. L. Mechanisms of alternative pre-messenger RNA splicing. Annual review of biochemistry 72, 291-336 (2003).

13 Hesselberth, J. R. Lives that introns lead after splicing. Wiley interdisciplinary reviews. RNA 4, 677-691, doi:10.1002/wrna.1187 (2013).

14 Spingola, M., Grate, L., Haussler, D. & Ares Jr, M. Genome-wide bioinformatic and molecular analysis of introns in Saccharomyces cerevisiae. Rna 5, 221-234 (1999).

15 Davis, C. A., Grate, L., Spingola, M. & Ares Jr, M. Test of intron predictions reveals novel splice sites, alternatively spliced mRNAs and new introns in meiotically regulated genes of yeast. Nucleic acids research 28, 1700-1706 (2000).

16 Juneau, K., Palm, C., Miranda, M. & Davis, R. W. High-density yeast-tiling array reveals previously undiscovered introns and extensive regulation of meiotic splicing. Proceedings of the National Academy of Sciences of the United States of America 104, 1522-1527, doi:10.1073/pnas.0610354104 (2007).

17 Zhang, Z., Hesselberth, J. R. & Fields, S. Genome-wide identification of spliced introns using a tiling microarray. Genome research 17, 503-509, doi:10.1101/gr.6049107 (2007).

18 Ng, R., Domdey, H., Larson, G., Rossi, J. & Abelson, J. A test for intron function in the yeast actin gene. Nature 314, 183-184 (1985).

19 Parenteau, J. et al. Deletion of many yeast introns reveals a minority of genes that require splicing for function. Molecular biology of the cell 19, 1932-1941 (2008).

20 Hooks, K. B., Naseeb, S., Parker, S., Griffiths-Jones, S. & Delneri, D. Novel intronic RNA structures contribute to maintenance of phenotype in Saccharomyces cerevisiae. Genetics 203, 1469-1481 (2016).

21 Parenteau, J. et al. Introns within ribosomal protein genes regulate the production and function of yeast ribosomes. Cell 147, 320-331, doi:10.1016/j.cell.2011.08.044 (2011).

22 Bonnet, A. et al. Introns Protect Eukaryotic Genomes from Transcription-Associated Genetic Instability. Molecular cell 67, 608-621 e606, doi:10.1016/j.molcel.2017.07.002 (2017).

23 Petfalski, E., Dandekar, T., Henry, Y. & Tollervey, D. Processing of the precursors to small nucleolar RNAs and rRNAs requires common components. Molecular and cellular biology 18, 1181-1189 (1998).

24 Qu, L.-H. et al. U24, a novel intron-encoded small nucleolar RNA with two 12 nt long, phylogenetically conserved complementarities to 28S rRNA. Nucleic acids research 23, 2669-2676 (1995).

25 Gray, J. V. et al. “Sleeping beauty”: quiescence in Saccharomyces cerevisiae. Microbiol Mol Biol Rev 68, 187-206, doi:10.1128/MMBR.68.2.187-206.2004 (2004).

26 Wu, T.-T., Su, Y.-H., Block, T. M. & Taylor, J. M. Evidence that two latency-associated transcripts of herpes simplex virus type 1 are nonlinear. Journal of virology 70, 5962-5967 (1996).

27 Qian, L., Vu, M. N., Carter, M. & Wilkinson, M. F. A spliced intron accumulates as a lariat in the nucleus of T cells. Nucleic acids research 20, 5345-5350 (1992).

28 Lorsch, J. R., Bartel, D. P. & Szostak, J. W. Reverse transcriptase reads through a 2′-5′ linkage and a 2′-thiphosphate in a template. Nucleic acids research 23, 2811-2814 (1995).

29 Fourmann, J. B. et al. Dissection of the factor requirements for spliceosome disassembly and the elucidation of its dissociation products using a purified splicing system. Genes & development 27, 413-428, doi:10.1101/gad.207779.112 (2013).

30 Martin, A., Schneider, S. & Schwer, B. Prp43 is an essential RNA-dependent ATPase required for release of lariat-intron from the spliceosome. Journal of Biological Chemistry 277, 17743-17750 (2002).

31 Katz, Y., Wang, E. T., Airoldi, E. M. & Burge, C. B. Analysis and design of RNA sequencing experiments for identifying isoform regulation. Nature methods 7, 1009-1015, doi:10.1038/nmeth.1528 (2010).

32 LaCava, J. et al. RNA degradation by the exosome is promoted by a nuclear polyadenylation complex. Cell 121, 713-724 (2005).

33 Jia, H. et al. The RNA helicase Mtr4p modulates polyadenylation in the TRAMP complex. Cell 145, 890-901, doi:10.1016/j.cell.2011.05.010 (2011).

34 Qin, D., Huang, L., Wlodaver, A., Andrade, J. & Staley, J. P. Sequencing of lariat termini in S. cerevisiae reveals 5′ splice sites, branch points, and novel splicing events. Rna 22, 237-253, doi:10.1261/rna.052829.115 (2016).

35 Terashima, H., Hamada, K. & Kitada, K. The localization change of Ybr078w/Ecm33, a yeast GPI-associated protein, from the plasma membrane to the cell wall, affecting the cellular function. FEMS microbiology letters 218, 175-180 (2003).

36 Loewith, R. & Hall, M. N. Target of rapamycin (TOR) in nutrient signaling and growth control. Genetics 189, 1177-1201 (2011).

37 Saxton, R. A. & Sabatini, D. M. mTOR signaling in growth, metabolism, and disease. Cell 168, 960-976 (2017).

38 Loewith, R. et al. Two TOR complexes, only one of which is rapamycin sensitive, have distinct roles in cell growth control. Molecular cell 10, 457-468 (2002).

39 Jorgensen, P. et al. A dynamic transcriptional network communicates growth potential to ribosome synthesis and critical cell size. Genes & development 18, 2491-2505 (2004).

40 Goranov, A. I. et al. Changes in cell morphology are coordinated with cell growth through the TORC1 pathway. Current Biology 23, 1269-1279 (2013).

41 Barnett, J. A. & Entian, K. D. A history of research on yeasts 9: regulation of sugar metabolism. Yeast 22, 835-894 (2005).

42 Zaragoza, D., Ghavidel, A., Heitman, J. & Schultz, M. C. Rapamycin induces the G0 program of transcriptional repression in yeast by interfering with the TOR signaling pathway. Molecular and cellular biology 18, 4463-4470 (1998).

43 Lempiäinen, H. et al. Sfp1 interaction with TORC1 and Mrs6 reveals feedback regulation on TOR signaling. Molecular cell 33, 704-716 (2009).

44 Mulleder, M. et al. Functional Metabolomics Describes the Yeast Biosynthetic Regulome. Cell 167, 553-565 e512, doi:10.1016/j.cell.2016.09.007 (2016).

45 Rousseau, A. & Bertolotti, A. An evolutionarily conserved pathway controls proteasome homeostasis. Nature 536, 184-189, doi:10.1038/nature18943 (2016).

46 Aronova, S., Wedaman, K., Anderson, S., Yates III, J. & Powers, T. Probing the membrane environment of the TOR kinases reveals functional interactions between TORC1, actin, and membrane trafficking in Saccharomyces cerevisiae. Molecular biology of the cell 18, 2779-2794 (2007).

47 Zurita-Martinez, S. A., Puria, R., Pan, X., Boeke, J. D. & Cardenas, M. E. Efficient Tor signaling requires a functional class C Vps protein complex in Saccharomyces cerevisiae. Genetics (2007).

48 Vyas, V. K. et al. New CRISPR Mutagenesis Strategies Reveal Variation in Repair Mechanisms among Fungi. mSphere 3, doi:10.1128/mSphere.00154-18 (2018).

49 Vyas, V. K., Barrasa, M. I. & Fink, G. R. A Candida albicans CRISPR system permits genetic engineering of essential genes and gene families. Science advances 1, e1500248 (2015).

50 Pleiss, J. A., Whitworth, G. B., Bergkessel, M. & Guthrie, C. Rapid, transcript-specific changes in splicing in response to environmental stress. Molecular cell 27, 928-937, doi:10.1016/j.molcel.2007.07.018 (2007).

51 Bergkessel, M., Whitworth, G. B. & Guthrie, C. Diverse environmental stresses elicit distinct responses at the level of pre-mRNA processing in yeast. Rna 17, 1461-1478, doi:10.1261/rna.2754011 (2011).

52 Munding, E. M., Shiue, L., Katzman, S., Donohue, J. P. & Ares Jr, M. Competition between pre-mRNAs for the splicing machinery drives global regulation of splicing. Molecular cell 51, 338-348, doi:10.1016/j.molcel.2013.06.012 (2013).

53 Warner, J. R. The economics of ribosome biosynthesis in yeast. Trends in biochemical sciences 24, 437-440 (1999).

54 Gasch, A. P. et al. Genomic expression programs in the response of yeast cells to environmental changes. Molecular biology of the cell 11, 4241-4257 (2000).

55 Chu, S. et al. The transcriptional program of sporulation in budding yeast. Science 282, 699-705 (1998).

56 Zheng, X.-F., Fiorentino, D., Chen, J., Crabtree, G. R. & Schreiber, S. L. TOR kinase domains are required for two distinct functions, only one of which is inhibited by rapamycin. Cell 82, 121-130 (1995).

## Online-only References

57 Weinberg, D. E. et al. Improved Ribosome-Footprint and mRNA Measurements Provide Insights into Dynamics and Regulation of Yeast Translation. Cell reports 14, 1787-1799, doi:10.1016/j.celrep.2016.01.043 (2016).

58 Harju, S., Fedosyuk, H. & Peterson, K. R. Rapid isolation of yeast genomic DNA: Bust n’Grab. BMC biotechnology 4, 8 (2004).

59 Subtelny, A. O., Eichhorn, S. W., Chen, G. R., Sive, H. & Bartel, D. P. Poly(A)-tail profiling reveals an embryonic switch in translational control. Nature 508, 66-71, doi:10.1038/nature13007 (2014).

60 Martin, M. Cutadapt removes adapter sequences from high-throughput sequencing reads. EMBnet. journal 17, pp. 10-12 (2011).

61 Dobin, A. et al. STAR: ultrafast universal RNA-seq aligner. Bioinformatics 29, 15-21 (2013).

62 Robinson, J. T. et al. Integrative genomics viewer. Nature biotechnology 29, 24-26 (2011).

63 Thorvaldsdóttir, H., Robinson, J. T. & Mesirov, J. P. Integrative Genomics Viewer (IGV): high-performance genomics data visualization and exploration. Briefings in bioinformatics 14, 178-192 (2013).

64 Bailey, T. L. et al. MEME SUITE: tools for motif discovery and searching. Nucleic acids research 37, W202-W208 (2009).

65 Wu, X. & Bartel, D. P. kpLogo: positional k-mer analysis reveals hidden specificity in biological sequences. Nucleic acids research (2017).

66 Pall, G. S., Codony-Servat, C., Byrne, J., Ritchie, L. & Hamilton, A. Carbodiimide-mediated cross-linking of RNA to nylon membranes improves the detection of siRNA, miRNA and piRNA by northern blot. Nucleic acids research 35, e60 (2007).

67 Chee, M. K. & Haase, S. B. New and Redesigned pRS Plasmid Shuttle Vectors for Genetic Manipulation of Saccharomycescerevisiae. G3 (Bethesda) 2, 515-526, doi:10.1534/g3.111.001917 (2012).

68 Gibson, D. G. et al. Enzymatic assembly of DNA molecules up to several hundred kilobases. Nature methods 6, 343-345 (2009).

69 Zalatan, J. G. et al. Engineering complex synthetic transcriptional programs with CRISPR RNA scaffolds. Cell 160, 339-350, doi:10.1016/j.cell.2014.11.052 (2015).

70 McFarlane, E. S. Ribonuclease activity during Gl arrest of the yeast Saccharomyces cerevisiae. Archives of microbiology 124, 243-247 (1980).

71 Wan, R., Yan, C., Bai, R., Lei, J. & Shi, Y. Structure of an intron lariat spliceosome from Saccharomyces cerevisiae. Cell 171, 120-132. e112 (2017).

